# Diverse haplotypes at a complex *Solanum americanum* locus confer resistance to *Phytophthora infestans* and *P. capsici*

**DOI:** 10.1101/2025.10.31.685743

**Authors:** Robert Heal, Andrea Carolina Olave-Achury, Maria Sindalovskaya, Li Long, Hari Sharan Karki, Jodie Taylor, Aleksandra Wawryk-Khamdavong, Kinga Bachowska, Matthew Smoker, Maheen Alam, Sarah Pottinger, Vatsal Arora, Kee-Hoon Sohn, Kamil Witek, Xiao Lin, Jonathan D G Jones

## Abstract

Plants encounter diverse pathogens and have evolved a two-layered innate immune system to detect pathogen molecules and activate defense mechanisms that restrict infection. Most cloned plant *Resistance* (*R*) genes encode NLR immune receptors. *NLR* genes are often found in clusters of paralogs with sequence and copy number variation; whether these *NLR* clusters evolve in response to single or multiple pathogens has been unclear. We report here the isolation of a *Phytophthora capsici* resistance gene, *Rpc2*, along with a novel *P. infestans* resistance gene, *Rpi-amr5*, from two *Solanum americanum* accessions. These orthologous genes reside in the *Rpi-amr1* cluster, which has previously been associated with resistance to *P. infestans*. By screening RXLR effector libraries of *P. infestans* and *P. capsici*, we identified multiple effectors recognized by both NLRs. Our findings highlight the complexity of *NLR* clusters and evolution driven by interactions with multiple pathogens. This work will underpin efforts to elevate resistance against *Phytophthora* pathogens and enhances our understanding of NLR evolution.

## Introduction

Oomycete species in the genus *Phytophthora* cause disease in many economically important plant species. Of these, the most notorious pathogen is *Phytophthora infestans*, the causal agent of potato late blight, which provoked the Irish famine in the 1840s and continues to cause significant damage to potato production (Savary et al., 2019, Fry, 2008). *P. capsici* is also a highly destructive oomycete pathogen with a broad host range including many important crops such as tomato, pepper, pumpkin, and melon (Kamoun et al., 2015). To date, nearly 50 Resistance to *Phytophthora infestans* (*Rpi*) genes have been cloned (Paluchowska et al., 2022), but little is known about *P. capsici* resistance.

Plant *Resistance* (*R*) genes often encode intracellular nucleotide-binding leucine-rich repeat (NLR) immune receptors, which recognise pathogen effectors and activate effector-triggered immunity (ETI). Most NLRs act as sensors to initiate an immune response after effector detection. While some of these are ‘functional singletons’, many NLRs require additional ‘helper’ NLRs to activate immunity (Wu et al., 2017, Witek et al., 2021, Lin et al., 2023, Lin et al., 2022, Ahn et al., 2023, Contreras et al., 2023b).

Most plant species resist most plant pathogens. This non-host resistance (NHR) is considered broader and more durable than host resistance (Lee et al., 2016, Vega-Arreguin et al., 2017, Zimnoch-Guzowska et al., 2003, Strugala et al., 2015, Panstruga and Moscou, 2020, Schulze-Lefert and Panstruga, 2011). *Solanum americanum* is a wild diploid species, which is considered a non-host to *P. infestans* (Colon et al., 1992) and is a useful model for studying plant-microbial interactions and plant immunity. We obtained 54 accessions from gene banks and generated reference-quality genomes for 4 accessions (Lin et al., 2023). By combining map-based cloning, SMRT-RenSeq, BSA-RenSeq, effectoromics, and GWAS, we cloned five NLR genes along with their corresponding recognized effectors, including Rpi-amr3/AVRamr3, Rpi-amr1/AVRamr1, Rpi-amr4/AVRamr4, R02860/PITG_02860, and R04373/ PITG_04373 (Lin et al., 2023, Lin et al., 2022, Lin et al., 2020, Witek et al., 2016, Witek et al., 2021). Remarkably, no *P. infestans* strains have been found to overcome either Rpi-amr1 or Rpi-amr3 (Lin et al., 2022, Witek et al., 2021). Understanding the durability of these non-host resistance genes will aid in effectively deploying resistance to ensure its longevity in crops.

Most characterized oomycete effector genes encode proteins with a signal peptide followed by both RXLR (Arg-X-Leu-Arg) and EER (Glu-Glu-Arg) motifs. The *P. infestans* T30-4 reference genome encodes 563 predicted RXLR effectors (Haas et al., 2009, Vleeshouwers et al., 2011). Some effectors lack RXLR or EER motifs but share with many RxLR effectors the presence of WY or LWY domains in the secreted protein (Bailey et al., 2011, Kim et al., 2024, Wood et al., 2020). This has enabled high-throughput effectoromics screening, and led to the identification of multiple *Rpi* and *Avr* genes (Haas et al., 2009, Vleeshouwers et al., 2011, Lin et al., 2023). However, no comprehensive *P. capsici* RXLR effector library has been reported.

Here, we describe the cloning and characterization of several novel *R* genes that map to the *Rpi-amr1* locus. *Rpi-amr5* was identified from a *P. infestans*-resistant accession lacking functional alleles of *Rpi-amr1* or *Rpi-amr3*. Interestingly, from a *P. capsici* resistant accession, we identified novel alleles of *Rpi-amr5* (named *Rpc2*), and *Rpi-amr1* (named *Rpi-amr1e-2308*), which confer resistance to both *P. capsici* and *P. infestans*. To investigate the molecular basis of *S. americanum* NHR to *P. infestans*, and its relationship to host resistance to *P. capsici*, we identified effectors recognized by these NLRs. Despite the difference in resistance to *P. capsici* conferred by *Rpi-amr5* and *Rpc2*, we found no difference in the recognition profile of the two NLRs. Rpi-amr5 and Rpc2 both recognise multiple effectors from *P. infestans* and *P. capsici*.

## Results

### *Rpi-amr5* from *S. americanum* accession SP2275 maps to the *Rpi-amr1* cluster

All tested *S. americanum* accessions resist *P. infestans* in the field, but five accessions show susceptibility in detached leaf assays (Lin et al., 2023). *Rpi-amr1* and *Rpi-amr3* were cloned from *S. americanum* accessions SP2273 and SP1102, respectively, and their cognate effectors, AVRamr1 and AVRamr3, were later defined. To identify novel *P. infestans* resistance, resistant *S. americanum* accessions were screened for a lack of response to these two effectors. The *S. americanum* accession SP2275 does not recognise AVRamr1, or AVRamr3, but is resistant to *P. infestans*.

SP2275 was crossed with a susceptible accession (SP2271) yielding resistant F1 plants. In an F2 population, phenotyping revealed a 3:1 (resistant: susceptible) (147 resistant and 53 susceptible; *χ*2 (1, N = 200) = 0.09, P = 0.6242) segregation ratio, indicating a single locus is responsible for resistance. To identify NLR-encoding genes linked to resistance in SP2275, 24 susceptible and 16 resistant segregants were used in an approach combining bulked segregant analysis (BSA) and resistance-gene enrichment sequencing (RenSeq) (Fig. 1a). Resistance mapped to the *Rpi-amr1* locus on the short arm of Chromosome 11 (Witek et al., 2021). However, SP2275 does not recognise AVRamr1 (Fig. 1b), therefore this resistance was named *Rpi-amr5*.

**Figure 1.**
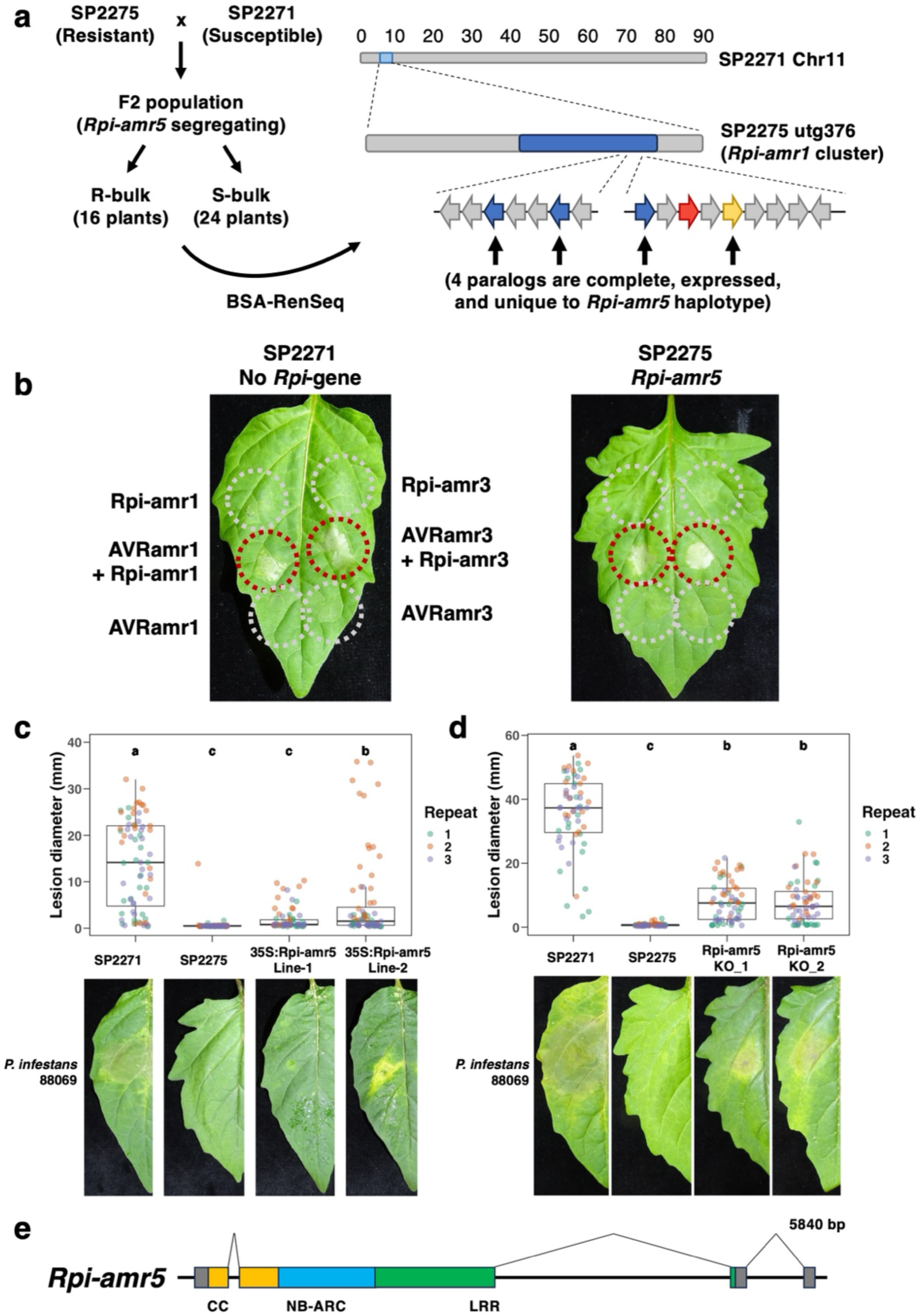
*Rpi-amr5* maps to the *Rpi-amr1* cluster and confers resistance to *P. infestans*. **(a)** SP2275 was crossed to SP2271, and an F2 population was phenotyped for foliar resistance to *P. infestans*. The 24 most susceptible and 16 most resistant plants were used for bulked segregant analysis using Resistance-gene enrichment sequencing (RenSeq). *Rpi-amr5* was positioned at the *Rpi-amr1* cluster on Chromosome 11, which contains 16 paralogs in SP2275. NLR-encoding genes are represented with arrows, *Rpi-amr5* candidates are indicated in blue. The *Rpi-amr1* ortholog (*Rpi-amr1e*, indicated in red) is non-functional in SP2275 due to a premature stop codon. The paralog corresponding to *Rpi-amr5* is indicated in yellow. **(b)** *S. americanum* accession SP2275 does not respond to AVRamr1 or AVRamr3. The effectors were transiently expressed in leaves of SP2275 and the susceptible reference accession SP2271. As a positive control for effector expression, the corresponding NLR was transiently co-expressed. *Agrobacterium* strains were infiltrated at an OD_600_ of 0.5; photographs were taken at 3 dpi. **(c)** Stable transformation with the functional *Rpi-amr5* elevates resistance to *P. infestans* in the susceptible accession SP2271. Leaves were inoculated with 10 μl drops of *P. infestans* zoospores (strain 88069), at a concentration of 20,000 spores mL^-1^ and imaged at 5 days post-inoculation. Plants derived from two independent primary transgenic events are shown. Three biological replicates were performed; all data points (72 per treatment) are represented as box-and-whisker plots. Statistical differences were determined using one-way ANOVA and Tukey’s HSD test (p < 0.05). **(d)** CRISPR knockout of *Rpi-amr5* in SP2275 reduces resistance to *P. infestans* relative to wild-type SP2275. Two independent biallelic *rpi-amr5* mutants were generated. Leaves were inoculated 10 μl droplets of *P. infestans* zoospores (20,000 spores mL^-1^) and imaged at 10 days post-inoculation. *P. infestans* strain 88069 was used. Three biological replicates were performed, all data points (60 per treatment) are represented as box-and-whisker plots. Statistical differences were determined using one-way ANOVA and Tukey’s HSD test (p < 0.05). **(e)** *Rpi-amr5* is a four-exon gene which encodes an 897 amino acid CC-NLR with 74.1% identity to Rpi-amr1. Intron and exon structure was determined using cDNA RenSeq data generated from SP2275 leaf tissue, no evidence of alternative splicing was found. The region encoding the CC domain is indicated in yellow, the NB-ARC in blue. and the LRR in green. The 5’ and 3’ UTRs are indicated in grey. The length of *Rpi-amr5* including UTRs, as predicted using cDNA RenSeq data, is shown.

To identify *Rpi-amr5*, we annotated the NLR-encoding genes within the *Rpi-amr1* cluster of SP2275. Using NLR-parser (Steuernagel et al., 2015), 16 paralogs within the *Rpi-amr1* cluster of SP2275 were identified. These paralogs form two sub-clusters: one was previously identified by Witek et al. (2021) and the other is more distal (Fig. 1a) (Supplementary Fig. 1). Gene models and expression levels of candidates were defined using cDNA RenSeq reads that were mapped to the SP2275 genome. Of the sixteen genes in the cluster, four are expressed and encode complete NLRs that have all canonical domains (Supplementary Table 1). These genes were named based on homology to NLRs in accession SP2273, where the *Rpi-amr1* cluster was first characterised (Witek et al., 2021) (Supplementary Fig. 1). Clear orthologs were identified for two candidates, *Rpi-amr1c-2275* and *Rpi-amr1h-2275*. However, two candidates lacked corresponding NLRs in SP2273 and were named based on their position within the SP2275 cluster: NLR_3-2275 and NLR_6-2275. The SP2273 *Rpi-amr1* gene that confers resistance to *P. infestans* is paralog *Rpi-amr1*e; however, its ortholog in SP2275 is non-functional due to a frameshift leading to a premature stop codon (Supplementary Fig. 2). This is consistent with SP2275’s absence of AVRamr1 response (Fig. 1b) and indicates that *Rpi-amr5* is a novel *Rpi* gene against *P. infestans*.

### VIGS and CRISPR define the paralog that confers *Rpi-amr5* function

To test the function of the four *Rpi-amr5* candidates, their open reading frames were amplified from SP2275 gDNA and cloned into expression vectors under the CaMV *35S* promoter. However, *Agrobacterium*-mediated transient expression in *Nicotiana benthamiana* resulted in constitutive, pathogen-independent cell death for two candidates, with weaker cell death observed for a third (Supplementary Fig. 3a). To assay candidates more effectively, TRV-based virus-induced gene silencing (VIGS) was used.

To silence the entire cluster, a fragment from the NB-ARC encoding region of the *Rpi-amr1h*-2275 paralog was cloned into a TRV2 vector, as this sequence is highly conserved among all members of this cluster. Individual paralogs were silenced by targeting sequence-diverged LRR-encoding regions (Supplementary Fig. 3b). Silencing of the *Rpi-amr1* family resulted in susceptibility, and silencing of *Rpi-amr1c-2275* produced a similar phenotype (Supplementary Fig. 3c), suggesting *Rpi-amr1c-2275* is likely to be the functional gene conferring resistance. This gene was subsequently named *Rpi-amr5*.

To validate its function, *Rpi-amr5* was stably transformed into the susceptible *S. americanum* accession SP2271, a background in which no constitutive activity was observed following transient expression of *Rpi-amr5* (Supplementary Fig. 4). Two independent transgenic lines showed elevated late blight resistance (Fig. 1c). Additionally, CRISPR/Cas9 mutagenesis was used to knock out *Rpi-amr5* in SP2275, leading to increased susceptibility relative to the wild-type accession (Fig. 1d, Supplementary Fig. 5). *Rpi-amr5* is a four-exon *Rpi-amr1* paralog which encodes a CC-NLR immune receptor (Fig 1e). The elevated susceptibility of *rpi-amr5* mutants confirms that this paralog is a major contributor to resistance. However, these mutants remain more resistant than the susceptible accession SP2271 (Fig 1d) which suggests additional genes within the same cluster may also contribute to resistance in SP2275.

### Screening *S. americanum* diversity for *P. capsici* resistance reveals *Rpc2* in SP2308

While *S. americanum* has been shown to carry resistance genes for *P. infestans*, *Pseudomonas syringae*, and *Ralstonia solanacearum* (Witek et al., 2016, Moon et al., 2021, Kim et al., 2025), resistance to *P. capsici* has not been previously explored. Screening of 52 *S. americanum* accessions revealed strong resistance in accession SP2308. This accession belongs to *S. americanum* group four, which shares characteristics with *Solanum nigrescens*, another diploid Solanum species. SP2308 was crossed with the susceptible accession SP2271. An F2 population of 185 plants segregated in a 3:1 (resistant: susceptible) ratio (140 resistant and 45 susceptible; *χ*2 (1, N = 185) = 0.01689, P = 0.83192), indicating that resistance is controlled by a single locus. This locus was named *Resistance to Phytophthora capsici 2* (*Rpc2*) (Supplementary Fig. 6a).

### Rpc2 maps to the Rpi-amr1/Rpi-amr5 cluster

To map *Rpc2*, short-read RenSeq was performed on bulked DNA from 34 susceptible and 30 resistant plants from the phenotyped F2 population. Bulked segregant analysis positioned *Rpc2* within the *Rpi-amr1*/*Rpi-amr5* cluster. Fine mapping using 78 susceptible segregants further narrowed the interval to 508 Kb (Supplementary Fig. 6a). Using PacBio RenSeq of SP2308, we identified five complete NLR-encoding genes unique to this accession, which were retained as *Rpc2* candidates. These genes – *Rpi-amr1a-2308*, *Rpi-amr1c-2308, Rpi-amr1e-2308, Rpi-amr1h-2308* and *Rpi-amr1l-2308* - were named following the nomenclature described previously (Supplementary Fig. 6b). The candidates include alleles of *Rpi-amr1* and *Rpi-amr5*. Subsequent testing was conducted to identify the functional *Rpc2* gene.

### *Rpc2* is an allele of *Rpi-amr5; Rpi-amr1e-2308* also confers resistance to *P. capsici*

Five *Rpc2* candidates were initially tested in *N. benthamiana* transient assays. Two of these NLRs, including the *Rpi-amr5* ortholog, conferred constitutive HR in *N. benthamiana* (Supplementary Fig. 7a). To further investigate *Rpc2*-mediated resistance, VIGS was used to test functionality of the candidates in the native accession. Silencing the SP2308 *Rpi-amr5* allele (*Rpi-amr1c-2308*) resulted in susceptibility to *P. capsici* (Supplementary Fig. 7b). Moreover, transforming *Rpi-amr1c-2308* into the susceptible *S. americanum* accession SP2271 conferred resistance to both *P. capsici* and *P. infestans* (Fig. 2a, Fig. 2b). These findings confirm that the primary source of resistance to *P. capsici* in SP2308 is *Rpi-amr1c-2308*, which we designate as *Rpc2*.

**Figure 2.**
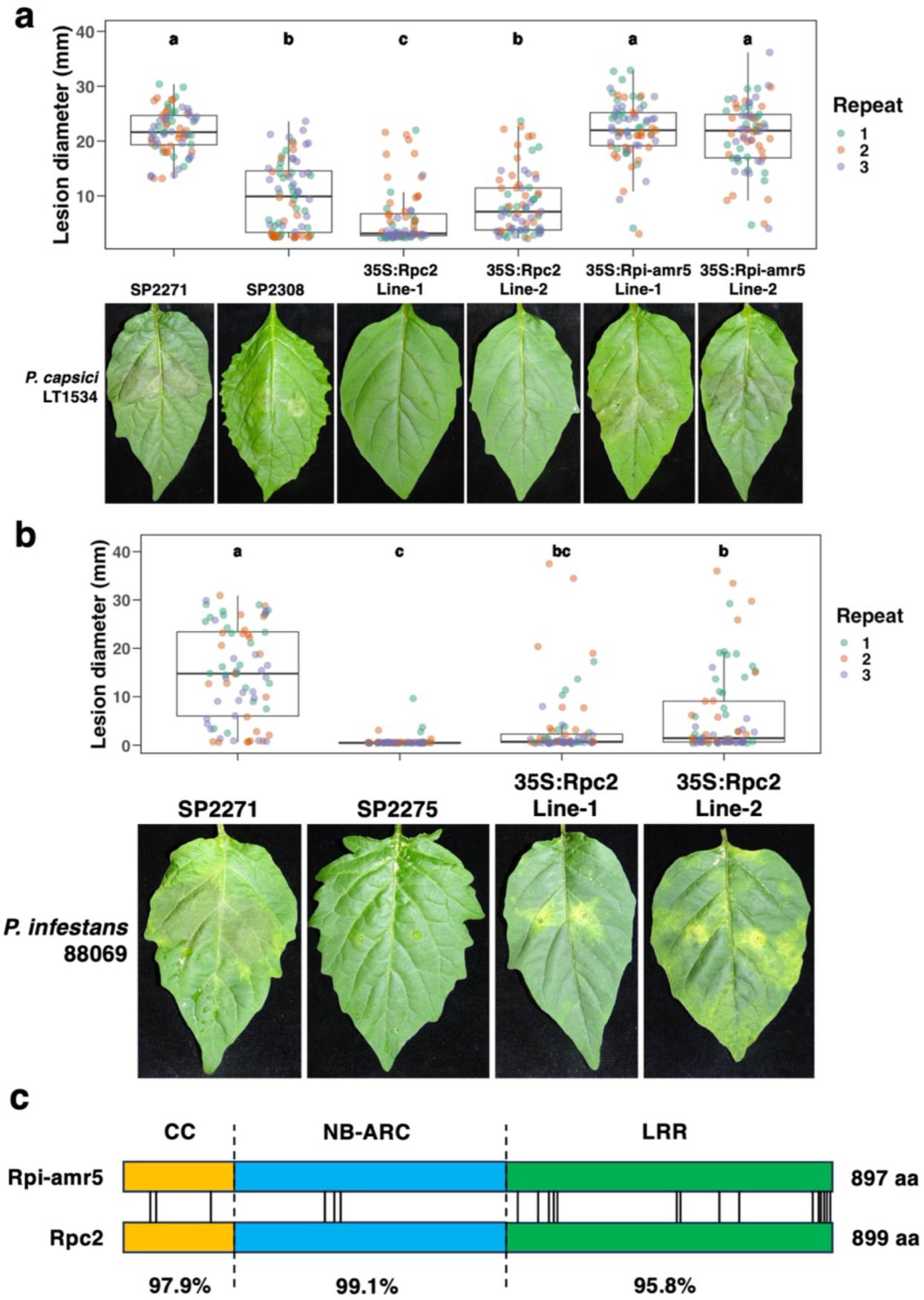
*Rpc2* is the *Rpi-amr5* allele in SP2308 and confers resistance to both *P. infestans* and *P. capsici* in *S. americanum.* **(a)** Overexpression of *Rpc2*, but not *Rpi-amr5*, elevates resistance to *P. capsici* in the susceptible *S. americanum* SP2271 background. Leaves were inoculated with *P. capsici* mycelial plugs (strain LT1534) and imaged at 3 days post-inoculation. Plants derived from two independent transgenic events are shown. Three biological replicates were performed, all data points (72 per treatment) are represented as box-and-whisker plots. Statistical differences were determined using one-way ANOVA and Tukey’s HSD test (p < 0.05). **(b)** Overexpression of *Rpc2* elevates resistance to *P. infestans* in the susceptible *S. americanum* SP2271 background. Leaves were inoculated with 10 μl drops of *P. infestans* strain 88069 (20,000 spores ml^-1^) and imaged at 3 days post-inoculation. Two independent transgenic events are shown. Three biological replicates were performed; all data points (72 per treatment) are represented as box-and-whisker plots. Statistical differences were determined using one-way ANOVA and Tukey’s HSD test (p < 0.05). **(c)** *Rpc2* and *Rpi-amr5* are alleles and have highly similar predicted amino acid sequences (97.4% identical). The positions of polymorphisms between the two predicted sequences are indicated.

*Rpc2* and *Rpi-amr5* share 97.4% amino acid identity (Fig. 2c). While both confer resistance to *P. infestans* in SP2308 and SP2275, respectively, only *Rpc2* is capable of conferring resistance to *P. capsici* (Fig. 2a) (Supplementary Fig. 8). We tested *Rpi-amr5* and *Rpc2* function in *Solanum lycopersicum*, which is susceptible to both *P. infestans* and *P. capsici*. The genes were recloned into a novel vector containing *SaNRC4a* promoter and terminator to ensure moderate expression levels (Supplementary Fig. 9a). These constructs were transformed into the susceptible tomato variety Moneymaker, and stable transformants were evaluated through *P. capsici* and *P. infestans* infection assays. Transformants expressing *Rpc2* were resistant to both pathogens, whereas *Rpi-amr5* transformants exhibited resistance only to *P. infestans* (Fig. 3a, Fig. 3b). These results suggest that the inability of *Rpi-amr5* to confer *P. capsici* resistance is likely due to allelic differences between *Rpi-amr5* and *Rpc2*, rather than their expression levels.

**Figure 3.**
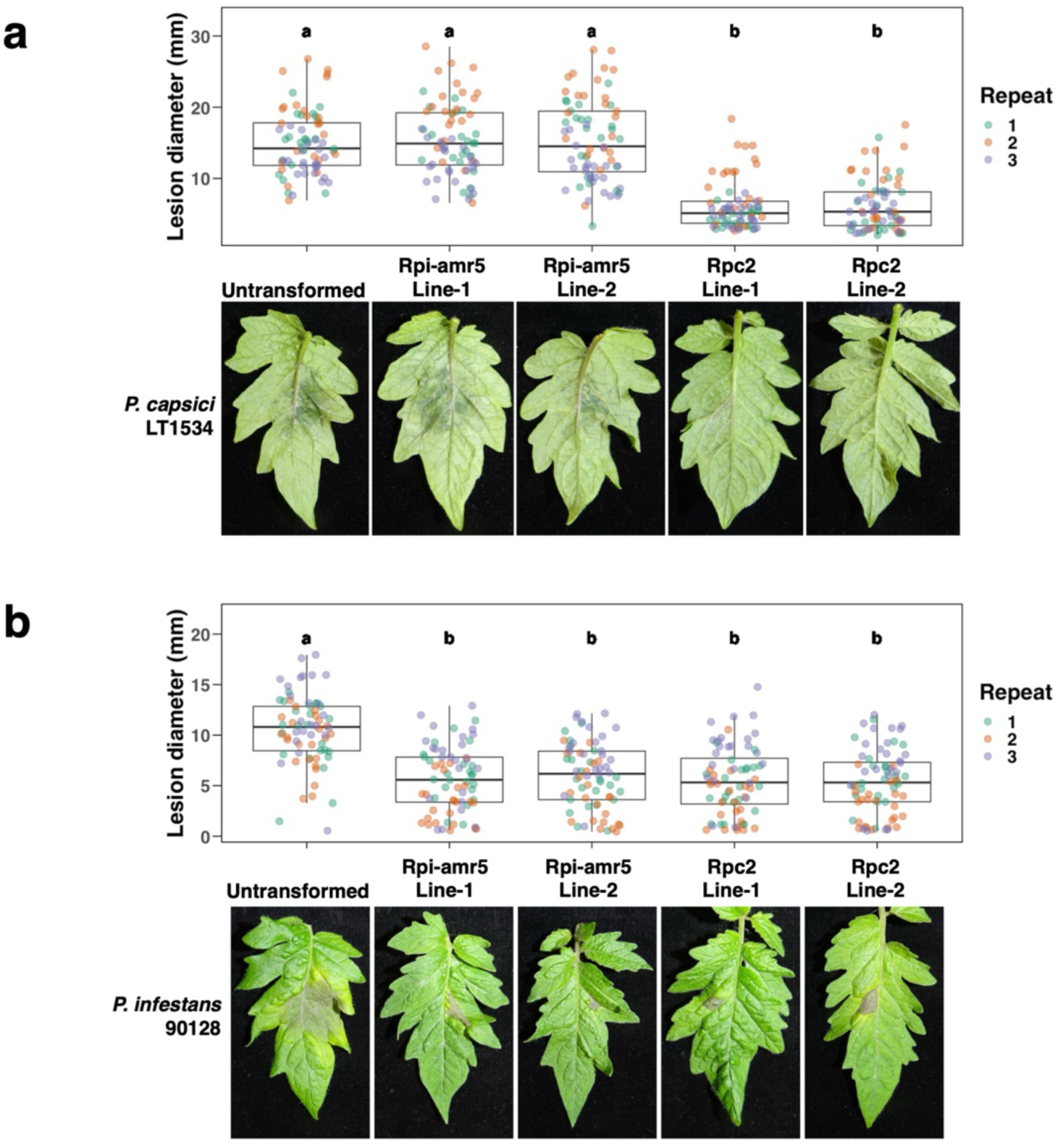
*Rpc2* and *Rpi-amr5* confer *P. infestans* resistance in transgenic *S. lycopersicum*; only *Rpc2* confers resistance to *P. capsici*. **(a)** *S. lycopersicum* (variety Moneymaker) lines expressing *Rpc2* under *SaNRC4a* regulatory elements are resistant to *P. capsici*, whereas *Rpi-amr5* transgenic lines show no increased resistance compared to the untransformed control. Two independent representative transgenic lines for each construct are shown. Leaves were inoculated with *P. capsici* (strain LT1534) mycelial plugs and imaged 3 days post-inoculation. **(b)** *S. lycopersicum* transgenic lines expressing either *Rpi-amr5* or *Rpc2* are resistant to *P. infestans*, while the untransformed control is susceptible. Leaves were inoculated with 10 μl drops of *P. infestans* strain 90128 (2,000 spores mL^-1^) and imaged at 5 dpi.

In SP2308, quantitative resistance to *P. capsici* was maintained even when *Rpc2* was mutated (Fig. 4a Supplementary Fig. 10), suggesting the presence of an additional resistance gene. To explore this, we tested three candidates which lacked constitutive activity in *N. benthamiana* (Supplementary Fig. 7a). Leaves expressing these genes were inoculated with *P. capsici* and pathogen growth assessed 72 hours post-inoculation. Notably, the SP2308 allele of the late blight resistance gene *Rpi-amr1e (Rpi-amr1e-2308*) significantly reduced *P. capsici* growth compared to *Rpi-amr3* or *Rpi-amr1e-2273*, both of which lack *P. capsici* resistance (Fig. 4b, Supplementary Fig. 11). Interestingly, both *Rpi-amr1e-2308* and *Rpi-amr1e-2273* recognise orthologs of AVRamr1 from *P. infestans* (PiAVRamr1) and *P. capsici* (PcapAVRamr1) (Fig. 4c) (Supplementary Fig. 12). This suggests that *Rpi-amr1e-2308* contributes to *P. capsici* resistance in SP2308.

**Figure 4.**
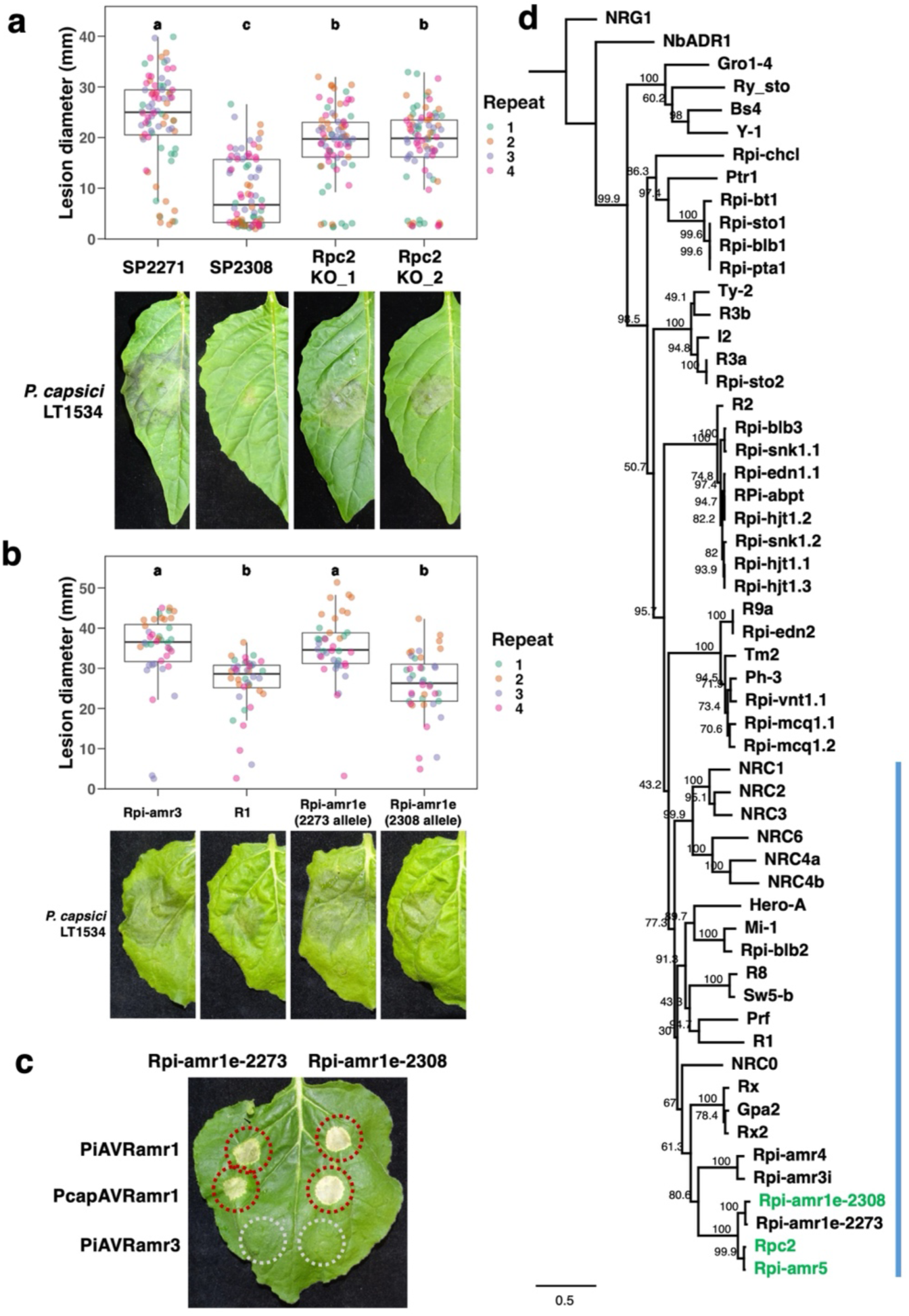
*Rpi-amr1e-2308* also confers resistance to *P. capsici.* **(a)** CRISPR-induced knockout of *Rpc2 (Rpi-amr1c)* in SP2308 reduces resistance to *P. capsici*, but some resistance is maintained relative to the susceptible control SP2271. Leaves were inoculated with *P. capsici* (strain LT1534) mycelial plugs and imaged 3 days post-inoculation. Two independent biallelic mutants are shown. Four biological replicates were performed; all data points (80 per treatment) are represented as box-and-whisker plots. Statistical differences were determined using one-way ANOVA and Tukey’s HSD test (p < 0.05). **(b)** *Rpi-amr1e-2308* confers partial resistance to *P. capsici* in *N. benthamiana* transient assays. *R1* was used as a positive control for *P. capsici* resistance. Agroinfiltration was performed using suspensions normalised to OD_600_ = 0.5. At 2 dpi, leaves were detached and *P. capsici* mycelial plugs were transferred onto them. Lesion size was measured 3 days post-inoculation. Four biological replicates were performed; all data points (40 per treatment) are represented as box- and-whisker plots. Statistical differences determined using one-way ANOVA and Tukey’s HSD test (p < 0.05). **(c)** Rpi-amr1-2308 and Rpi-amr1-2273 (with 92.3% amino acid identity) both confer recognition of AVRamr1 orthologs from *P. infestans* (PiAVRamr1) and *P. capsici* (PcapAVRamr1). *N. benthamiana* leaves were infiltrated with *Agrobacterium* suspensions at OD_600_ = 0.2; two days later leaves were inoculated. Leaves were imaged at 4 dpi. **(d)** Phylogenetic tree of Rpi-amr5, Rpc2, Rpi-amr1e-2308, and a panel of previously characterised Solanaceae NLRs. These three sensor NLRs are closely related to Rpi-amr1e-2273 and reside within the NRC superclade (indicated by the blue line). The tree was constructed using NB-ARC sequences. Branch support values are shown as percentages, based on 1000 iterations.

### Transient expression of *Rpc2* and *Rpi-amr5* with *S. americanum NRC2* enables identification of two recognised *P. infestans* effectors

Transient expression of *Rpi-amr5* or *Rpc2* under the CaMV *35S* or *SaNRC4a* promoters in *N. benthamiana* results in cell death (Supplementary Fig. 9b). To investigate this effector-independent cell death and enable effector screening, the role of NRC helper NLRs was investigated. Like Rpi-amr1; Rpi-amr5 and Rpc2 are within a branch of the NRC-dependent CC-NLRs (Fig. 4d). NRC proteins are helper NLRs through which cell death is activated after pathogen detection by sensor CC-NLRs (Contreras et al., 2023b, Wu et al., 2017). In *nrc2/3/4* mutant *N. benthamiana* plants (Wu et al., 2019), *Rpi-amr5* expression does not trigger cell death. Complementation with *35S:NbNRC2,* but not *35S:SaNRC2* (SaNRC2 = *S. americanum* NRC2), restored strong constitutive activity (Fig. 5a). Additional *NRC* genes cloned from *N. benthamiana* and *S. americanum* were co-expressed with *Rpi-amr5*. Unexpectedly, *Rpi-amr5* also displayed constitutive activity with *NbNRC3* and *SaNRC3* (Supplementary Fig. 14). While the underlying biology is unclear, this finding allowed the coexpression of *35S:SaNRC2* with *Rpc2* or *Rpi-amr5* in *nrc2/3/4* plants for effector screening.

**Figure 5.**
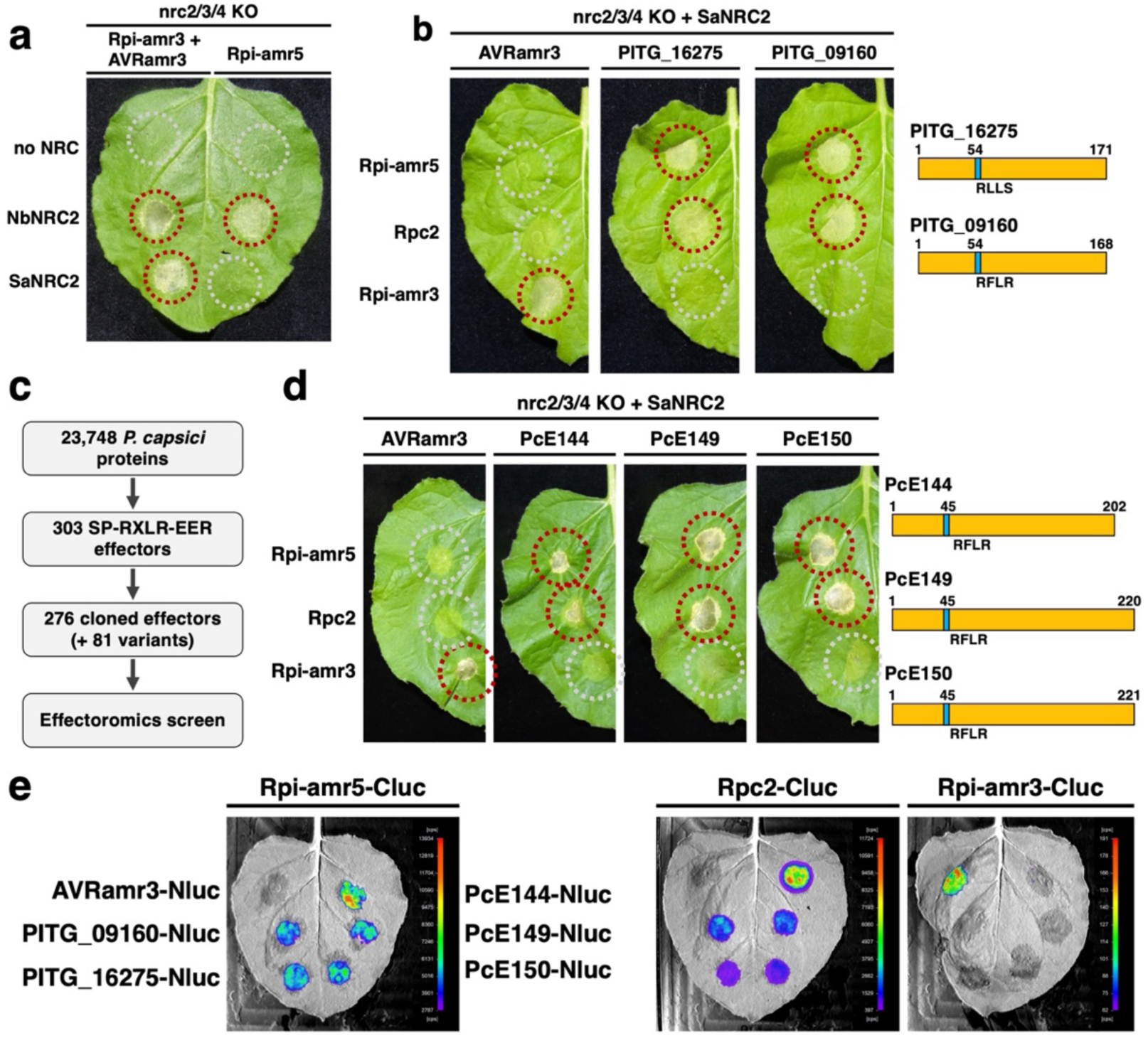
Rpi-amr5 and Rpc2 recognise two *P. infestans* effectors and at least three *P. capsici* effectors. **(a)** Absence of NRC helper NLRs abolishes Rpi-amr5/Rpc2 constitutive activity in *N. benthamiana*. Complementation with NbNRC2, but not SaNRC2, restores constitutive activity. *Agrobacterium* strains were infiltrated at OD_600_ = 0.2, and leaves were imaged at 3 dpi. **(b)** Rpi-amr5 and Rpc2 recognise PITG_09160 and PITG_16275 from *P. infestans*. Co-expression of *SaNRC2* with either allele, and either *PITG_09160* or *PITG_16275* results in effector-dependent HR. Expression of these effectors with *Rpi-amr3* and *SaNRC2* does not result in cell death. *Agrobacterium* strains were infiltrated at OD_600_ = 0.2, and leaves were imaged at 3 dpi. **(c)** *P. capsici* effector identification pipeline. From the predicted *P. capsici* proteome, SP-RXLR-EER effectors were identified by filtering for the presence of a signal peptide, RXLR motif, and EER motif. Of the 303 predicted effectors, 276 were cloned. During cloning, 81 additional allelic variants were identified. **(d)** Rpi-amr5 and Rpc2 both recognise at least three *P. capsici* effectors: PcE144, PcE149, and PcE150. Expression of these effectors with *SaNRC2* and *Rpi-amr5* induces cell death. A similar response was observed with Rpc2. No HR was detected when PcE144, PcE149, or PcE150 were co-expressed with Rpi-amr3. These three effectors are typical RXLR-EER-type effectors. All *Agrobacterium* strains were infiltrated at OD_600_ = 0.2. Leaves were imaged at 3 dpi. **(e)** A split luciferase assay was used to test for *in planta* association between the five recognized effectors (PITG_09160, PITG_16275, PcE144, PcE149, and PcE150) and both Rpi-amr5 and Rpc2. Luminescence was observed when Rpi-amr5, or Rpc2, but not Rpi-amr3, was co-expressed with the effectors. To avoid cell death, the assay was performed in the *nrc2/3/4* KO *N. benthamiana* line. Sensor NLRs were tagged with a C-terminal luciferase fragment, and effectors with an N-terminal luciferase fragment.

Given the near-identity of Rpi-amr5 and Rpc2, we hypothesised that they recognise the same *P. infestans* effector. All available *S. americanum* accessions have been screened for recognition of 315 *P. infestans* effectors (Lin et al., 2023). Two effectors were recognised by both SP2275 and SP2308, but not SP2271. When transiently expressed in *nrc2/3/4 N. benthamiana* with *35S:SaNRC2* and either *Rpi-amr5* or *Rpc2*, the effector PITG_16275 was recognised by both NLRs (Fig. 5b). Recognition of PITG_16275 by Rpi-amr5 has been independently verified in SP2297, another *S. americanum* accession. Using an F2 population, recognition was mapped to the same locus which contains an *Rpi-amr5* allele encoding an identical NLR (Supplementary Fig. 15). Effectoromics screening (Lin et al., 2023) showed that 24 of the 52 *S. americanum* accessions respond to PITG_16275, suggesting that functional *Rpi-amr5* alleles are widespread in *S. americanum*. Of the 24 accessions that respond to PITG_16275, 19 also respond to AVRamr1. Thus, in these accessions, *Rpi-amr1* and *Rpi-amr5* are clustered on the same haplotype and form a natural “stack” of *Rpi* genes.

A BLAST sequence homology search for similar *P. infestans* effectors revealed PITG_09160, which shares 39.2% amino acid identity with PITG_16275 after the EER motif. Three additional effectors - PITG_06076, PITG_06059 and PITG_06074 - were also identified, but these three effectors are not expressed during infection (Lin et al., 2020). Transient expression assays confirmed that *Rpi-amr5* and *Rpc2* recognise both PITG_09160 and PITG_16275, to elicit HR (Fig. 5b). After identifying the *P. infestans* elicitor of Rpi-amr5, we tested the dependence of this sensor on SaNRC proteins. In addition to SaNRC2, Rpi-amr5 can be supported by SaNRC1. This contrasts with Rpi-amr1 which is not supported by SaNRC1 (Supplementary Fig. 16) (Lin et al., 2023).

### Homologs of PITG_09160 and PITG_16275 are found in several *Phytophthora* species, but not *P. capsici*

*Rpi-amr5* and *Rpc2* recognise two *P. infestans* effectors, which share similar sequences. We hypothesised that *P. capsici* may contain a similar effector which is recognised by Rpc2. A BLAST search against both the *P. capsici* genome and predicted proteome (Stajich et al., 2021) found no proteins with homology to PITG_16275 or PITG_09160. Therefore, a structural homology search was performed using the bioinformatic tool Foldseek (van Kempen et al., 2024). From the *P. capsici* proteome prediction, 356 effectors were predicted based on the presence of a signal peptide, as well as EER and RXLR motifs (Fig. 5c). Alphafold prediction of these proteins was performed to assemble a library of predicted structures to use as a Foldseek database. However, no proteins were predicted to share high structural similarity with the recognised *P. infestans* proteins (Supplementary Fig. 17).

While no PITG_16275 homologs were identified in *P. capsici*, *P. palmivora*, *P. parasitica*, *P. megakarya* and *P. cactorum* were found to contain highly similar effectors (Supplementary Fig. 17). A high-scoring hit from each species was synthesised and transiently expressed with *Rpi-amr5*. Co-expression of the *P. megakarya*, *P. palmivora* and *P. parasitica* orthologs with *Rpi-amr5* caused strong cell death. For the *P. cactorum* ortholog, no clear HR was observed (Supplementary Fig. 18). While resistance to these species has not been tested, this experiment demonstrates that Rpi-amr5 can recognise effectors from at least five *Phytophthora* species. Interestingly, many more PITG_16275 homologs were predicted in *P. palmivora* and *P. megakarya* (21 and 28) than in *P. parasitica*, *P. cactorum,* or *P. infestans* (8, 2, and 5), suggesting an expansion of this effector group in these species. Although it remains unclear how many of these homologs are recognised by Rpi-amr5, in combination with the appropriate NRC, this sensor may prove to be a useful resistance gene against these species (Du et al., 2025).

### Three *P. capsici* effectors are recognised by both *Rpi-amr5* and *Rpc2*

No *P. capsici* effectors with homology to PITG_16275 or PITG_09160 were identified using sequence or structural homology searches. Therefore, we cloned a library of 277 FLAG-tagged *P. capsici* RxLR effectors (and 80 additional allelic variants) using a combination of gene synthesis and PCR amplification from *P. capsici* gDNA (Fig. 5c). Effectors were screened by transient expression in SP2308, SP2275 and SP2271. Effectors that elicit HR in SP2308 were then tested for recognition by *Rpc2* and *Rpi-amr5* in *N. benthamiana*. Three *P. capsici* effectors are recognised by Rpc2: PcE144, PcE149 and PcE150. Surprisingly, Rpi-amr5 also showed recognition of these effectors, despite not being able to confer *P. capsici* resistance in *S. americanum* or *S. lycopersicum* (Fig. 5d).

The three recognised *P. capsici* effectors have high sequence similarity to each other (65.4-92.5% amino acid identity), but low identity to the recognised *P. infestans* effectors (12.2-15.0% amino acid identity) (Supplementary Fig. 19). No orthologs of PcE144, PcE149 or PcE150 were identified in *P. infestans*, *P. parasitica*, *P. palmivora*, *P. megakarya* or *P. cactorum*.

### Rpi-amr5 and Rpc2 associate with their cognate effectors *in planta*

Rpi-amr5 and Rpc2 both recognise five effectors: two from *P. infestans* and three from *P. capsici*. These effectors vary significantly in length and amino acid sequence (Fig. 5b, Fig. 5d, Supplementary Fig. 19). To determine whether recognition involves direct association with the NLRs, a split luciferase assay was conducted to test for *in planta* association. In this assay, the NLRs were fused with the C-terminal fragment of luciferase, while effectors were tagged with the N-terminal fragment. Constructs were expressed in the *nrc2/3/4 N. benthamiana* line, to prevent cell death being triggered. Luminescence was observed when any of the five recognised effectors were co-expressed with either Rpi-amr5 or Rpc*2*, indicating direct association. In contrast, no luminescence was detected upon co-expression with Rpi-amr3, or the control effector AVRamr3, which does not activate Rpi-amr5 or Rpc2 (Fig. 5e). These results suggest that, despite the sequence diversity among the recognised effectors, both Rpi-amr5 and Rpc2 are activated through direct association. This finding supports a model of recognition based on physical proximity to the NLRs, rather than the modification of a host target or an indirect recognition mechanism.

### Solanum immune receptors convergently evolved recognition of Rpi-amr5/Rpc2 AVRs

Rpi-amr5 and Rpc2 recognise the *P. infestans* effectors PITG_09160 and PITG_16275. Notably, additional non-Rpi-amr5 recognition of PITG_09160 was observed in *S. americanum*. Screening *Rpi-amr5* and *Rpc2* knock-out lines for loss of recognition of the five known effectors revealed that SP2275 *rpi-amr5* mutants lost response to all five effectors. However, SP2308 *rpc2* lines retained recognition of PITG_09160, as well as PcE149 and PcE150 (Supplementary Fig. 20). This indicates the presence of additional receptors in SP2308 capable of detecting these effectors. Furthermore, other *S. americanum* accessions lacking functional *Rpi-amr5* alleles also respond to PITG_09160 (Supplementary Fig. 20). These findings suggest that immune receptors within *S. americanum* have independently evolved distinct mechanisms to recognise PITG_09160, demonstrating convergent evolution of immune receptor functionality within the species.

## Discussion

### Several distinct paralogs of the *Rpi-amr1* cluster confer *Phytophthora* resistance

The *Phytophthora* genus contains a diverse range of plant pathogens, many of which have significant economic impact. *P. infestans*, the causal agent of late blight, remains a global threat to potato production, while *P. capsici* causes disease in numerous *Solanum, Capsicum,* and Cucurbitaceae species. While *S. americanum* is a non-host to *P. infestans*, most *S. americanum* accessions are susceptible to *P. capsici*.

We show here that the *S. americanum* accession SP2275 carries *Rpi-amr5,* which encodes a CC-NLR immune receptor and confers resistance to *P. infestans* (Fig. 6a). In parallel, resistance to *P. capsici* was characterised in *S. americanum* accession SP2308. Two NLR-encoding genes located at the SP2308 *Rpi-amr1* locus confer resistance to both *P. capsici* and *P. infestans*; *Rpc2,* an allele of *Rpi-amr5*, confers most of the *P. capsici* resistance, while weaker resistance is conferred by *Rpi-amr1e-*2308, a novel allele of *Rpi-amr1* (Fig. 6a). Similarly, of 24 *Rpi-amr5*-containing haplotypes, 19 also carry a functional *Rpi-amr1* allele. These examples illustrate that gene stacks have often been selected for in nature.

**Figure 6.**
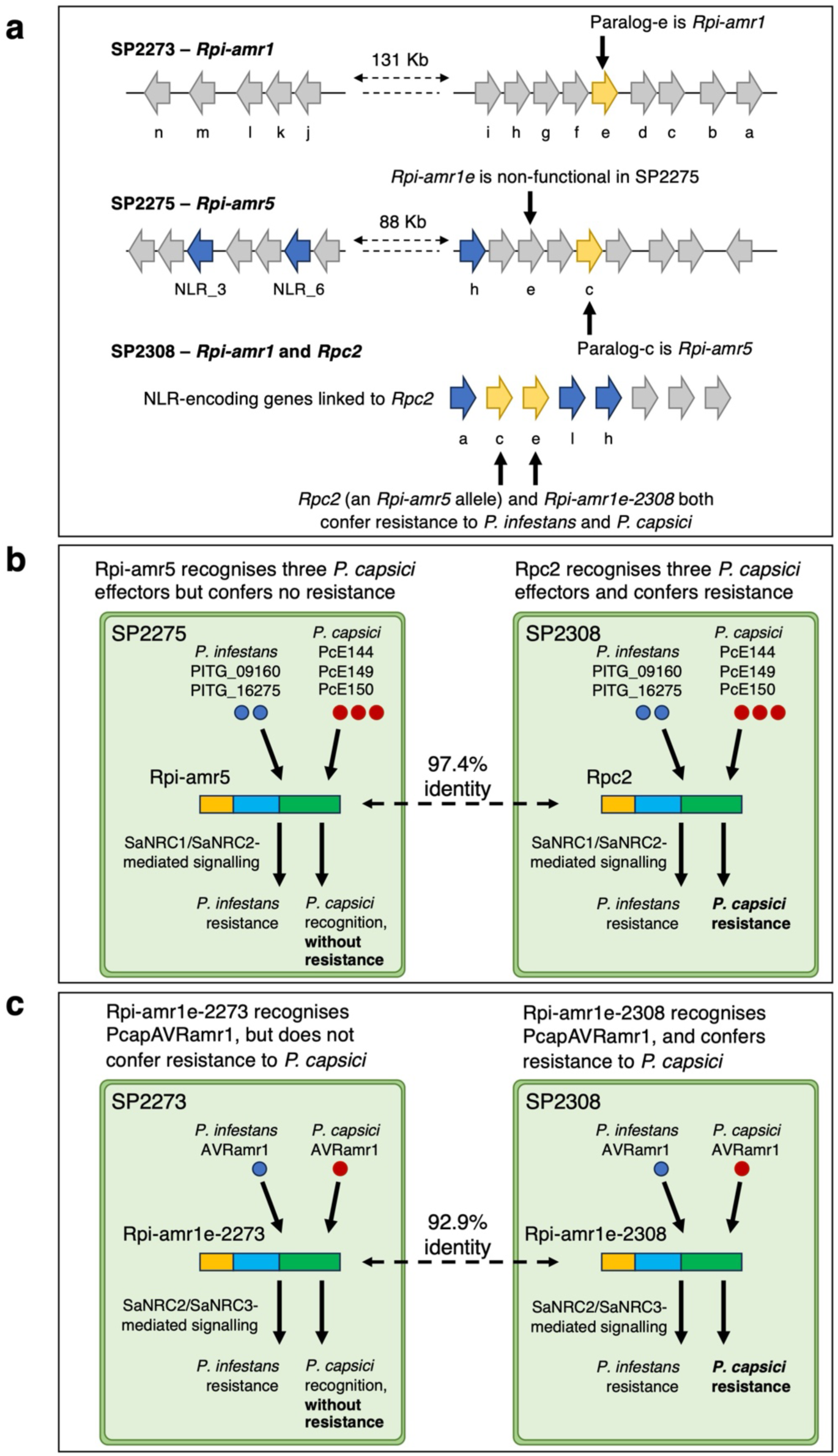
A complex *Solanum americanum* resistance gene locus confers resistance to *Phytophthora infestans* and *P. capsici*. **(a)** The *Rpi-amr1* clusters in *S. americanum* accessions SP2273 (the source of *Rpi-amr1e-2273*), SP2275 (the source of *Rpi-amr5*), and SP2308 (the source of *Rpc2* and *Rpi-amr1e-2308*) contain distinct genes that confer resistance to *P. infestans* and *P. capsici*. NLR-encoding genes are represented with arrows, genes known to confer resistance to either *P. infestans* or *P. capsici* are indicated in yellow. **(b)** *Rpi-amr5* and *Rpc2* were cloned from accessions SP2275 and SP2308, respectively. Rpc2 recognises two effectors from *P. infestans* and three from *P. capsici*, and confers resistance to both pathogens. *Rpi-amr5* (an allele of *Rpc2*) recognises the same effectors but does not confer resistance to *P. capsici*. **(c)** Rpi-amr1e-2273 recognises AVRamr1 orthologs from both *P. infestans* and *P. capsici*, but does not confer resistance to *P. capsici*. A novel allele, *Rpi-amr1e-2308*, confers both recognition and resistance.

### Interspecific combinations of helper NLR and sensor NLR can result in effector-independent defense activation

Both Rpi-amr5 and Rpc2 activate constitutive HR in *N. benthamiana*, but not in *nrc2/3/4* knockout lines. This constitutive activity can be restored in *nrc2/3/4* lines by co-expression with *35S:NbNRC2*, but not *35S:SaNRC2*, suggesting that inappropriate combinations of sensor and helper NLRs can be maladaptive. Recent studies have shown that the restricted taxonomic functionality (RTF) of NRC-dependent sensor NLRs results from the absence of a compatible helper NLR (Du et al., 2025, Goh et al., 2024). Our data suggest that in addition, interspecific combinations of sensor NLRs and NRCs can result in aberrant, effector-independent activation of helper NLRs of one species by sensor NLRs from another species. Conceivably, such outcomes could have arisen during introgression of *R* genes from wild relatives of crops, though deleterious combinations would have been selected against. *Rpc2* and *Rpi-amr1e-2308* may have broad utility, either through direct transfer into solanaceous hosts of *P. capsici*, such as tomato and pepper, or by transfer to less related species, such as cucurbits. Cucurbitaceae lack NRC proteins (Wu et al., 2017); however, co-transfer of *Rpc2* and *SaNRC2* into cucurbits could elevate resistance to *P. capsici* (Du et al., 2025).

### Rpi-amr5 and Rpc2 recognise at least five *Phytophthora* effectors

By co-expressing *Rpi-amr5* or *Rpc2* in *nrc2/3/4* lines together with *35S:SaNRC2* and clones from a *P. infestans* effector library, we identified PITG_16275 and PITG_09160 as *P. infestans* effectors recognized by Rpi-amr5 and Rpc2. However, no orthologs of PITG_16275 were identified in *P. capsici*. To identify *P. capsici* effectors recognised by Rpc2, we generated and screened a library of RXLR effectors in *S. americanum*. We identified three *P. capsici* effectors recognised by both Rpi-amr5 and Rpc2. Split luciferase assays suggest that the interaction between Rpi-amr5/Rpc2 and these five effectors is likely direct.

Many accessions lacking functional *Rpi-amr5* alleles nevertheless respond to PITG_09160. While response to PITG_09160 is lost in SP2275 *rpi-amr5* mutant lines, it is retained in SP2308 *rpc2* mutant lines. These data suggest that an additional PITG_09160 receptor is present in *S. americanum*. Recognition of PITG_09160 and PITG_16275 has also been reported in other Solanaceae species, indicating convergent evolution of receptors to recognise these effectors. Similar examples of convergent evolution include the recognition of AVR2, a *P. infestans* effector detected by both R2 and Rpi-mcq1, which were cloned from *Solanum demissum* and *Solanum mochiquense,* respectively (Aguilera-Galvez et al., 2018).

### Why does *Rpi-amr5* not confer resistance to *P. capsici*?

Three *P. capsici* effectors are recognized by Rpc2, and unexpectedly, Rpi-amr5 also recognises all these effectors. Despite this, *Rpi-amr5* does not confer resistance to *P. capsici* (Fig. 6b). Similarly, *Rpi-amr1e-2308* provides resistance to *P. capsici*, whereas the *Rpi-amr1e-2273* allele does not, despite both recognising the *P. capsici* AVRamr1 ortholog (Fig. 6c). This is consistent with previous reports of incomplete correspondence between HR to recognized effectors and disease resistance (Lin et al., 2023, Oh et al., 2024, Gilroy et al., 2011). Notably, *S. americanum* R02860 and R04373 recognise *P. infestans* effectors PITG_02860 and PITG_04373, but neither confer resistance (Lin et al., 2023). Although puzzling, this phenomenon is not unprecedented.

To understand the molecular basis for Rpc2-dependent resistance, several hypotheses warrant investigation. Conceivably, Rpc2 may recognise an additional effector that is not detected by Rpi-amr5. This effector might be absent from our library, which contains only RXLR-type effectors. Non-RXLR oomycete effectors are well-documented. For instance, *Hyaloperonospora arabidopsidis* effectors ATR2 and ATR5, which are recognised by *A. thaliana* NLRs RPP2A/RPP2B and RPP5, respectively, lack RXLR motifs but retain canonical effector features such as signal peptides, EER motifs, and WY/LWY domains (Bailey et al., 2011, Kim et al., 2024). Wood et al. (2020) reported that several oomycetes, including *P. capsici*, secrete proteins lacking RXLR motifs but containing EER motifs or WY domains. If such a non-RXLR effector is recognised by Rpc2 but not Rpi-amr5, it would explain the observed resistance differences. This explanation would be consistent with the fact that there are 19 polymorphisms in the LRRs compared to only 3 in the CC and 3 in the NB-ARC domains.

Another possible explanation involves allele-specific suppression of Rpi-amr5. Suppression of NLR-dependent immunity is a phenomenon reported in multiple studies (Chen et al., 2012, Yin et al., 2017, Derevnina et al., 2021, Contreras et al., 2023a). This suppression could explain why recognition does not always translate into resistance. Understanding the mechanisms underlying NLR suppression by effectors could pave the way for engineering resistant alleles to evade this suppression, thereby enhancing resistance (Contreras et al., 2023a).

### Host and non-host resistance in *S. americanum* directly overlap

Non-host resistance is a multifaceted phenomenon, often involving multiple mechanisms. As highlighted by Panstruga and Moscou (2020), NLR-mediated immunity contributes significantly to both host and non-host resistance. While *S. americanum* is a non-host to *P. infestans* in the field, resistance to *P. capsici* is uncommon, displaying features of host resistance. We show here that resistance to these two pathogens involves different haplotypes at the *Rpi-amr1* locus. Two paralogs - *Rpi-amr1* and *Rpi-amr5* - confer resistance to *P. infestans* in both haplotypes. In the *P. capsici* resistant accession SP2308, novel alleles of *Rpi-amr1* (*Rpi-amr1e-2308*) and *Rpi-amr5* (*Rpc2*) also provide resistance to *P. capsici*.

In characterising NLR-mediated non-host resistance to *P. infestans* in *S. americanum*, we have so far identified four *Rpi* genes that confer qualitative resistance: *Rpi-amr1*, *Rpi-amr3, Rpi-amr4,* and *Rpi-amr5.* All *S. americanum* accessions with qualitative resistance to *P. infestans* in detached leaf assays, as described by (Lin et al., 2023), contain at least one of these *Rpi* genes. The broad-spectrum resistance profiles of *Rpi-amr1* and *Rpi-amr3* suggest that they may derive durability from the presence of multiple *Rpi* genes. Further elucidation of the genetic and molecular basis of non-host resistance will aid in deploying resistance strategies that are both sustainable and durable in real-world conditions.

## Methods

### Constructs for transient and stable transformation

To verify candidate genes, ORFs were amplified and cloned into vectors containing the CaMV 35S promoter and the *Agrobacterium tumefaciens* Ocs terminator, using either GoldenGate or USER cloning as previously described (Witek et al., 2016, Lin et al., 2023). Candidates cloned using GoldenGate were ligated into pICLS86922, while USER clones were generated by ligation into pICSLUS0004OD. Additional constructs were generated to test *Rpi-amr5* and *Rpc2* under *S. americanum NRC4a* (*SaNRC4a*) regulatory elements. A custom vector (pICSLUS0005OD) was generated to accept each ORF following the USER method (Nour-Eldin et al., 2010). This vector includes a 943 bp promoter and a 793 bp terminator, both cloned from *S. americanum* accession SP1102.

*Agrobacterium* suspensions were prepared to OD described in the figure legends. For HR assays in *N. benthamiana*, and *S. americanum,* fully expanded leaves were selected from plants aged between five and six weeks. Leaves were spot-infiltrated using 1 ml needleless syringes. When multiple infiltrations were made on a single leaf, care was taken to avoid overlapping infiltration zones. Plants were left for 3-5 days to allow cell death to develop. To test candidate *Rpi* and *Rpc* genes, infiltration of whole leaves was performed. Leaves were left for 24 hours before being removed and inoculated as described below.

### Virus-induced gene silencing (VIGS) of *S. americanum*

Virus-induced gene silencing (VIGS) was used to silence candidate *R*-genes in *S. americanum*. A BsaI-compatible TRV2 vector was used (Duggan et al., 2021). To silence the *Rpi-amr1*/*Rpi-amr5*/*Rpc2* cluster, a highly conserved 265 bp fragment from the NB-ARC-encoding region of *Rpi-amr1h-2275*. To target individual paralogs, the least conserved regions were identified. These regions were generally shorter (between 51 and 115 bp). For each paralog, two separate fragments were used. A conservative approach was used to define off-target silencing: the candidate region was divided into 21-mer sequences, off-target hits were defined as those where fewer than 2 SNPs were present within these 21-mer sequences (i.e. ≥90.5% identity).

VIGS of *S. americanum* was performed by vacuum infiltration of seedlings between 2-5 days post-germination. *Agrobacterium* suspensions of TRV1 and TRV2 (with the silencing fragment) were mixed in a 1:1 ratio to a final OD_600_ of 0.3. Before infiltration, acetosyringone (500 µM) and Silwet-L77 (0.1 µl/ ml) were added. Seedlings were submerged into the *Agrobacterium* suspension, and a vacuum was applied for one minute, before being released. This was repeated two additional times. Following infiltration, seedlings were transferred to compost and grown in glasshouse conditions. When plants were five weeks old, leaves were selected and drop-inoculated with either *P. infestans* or *P. capsici* zoospores. To confirm the effectiveness of gene silencing, a fragment from the gene encoding the magnesium-chelatase subunit ChIH was used. Silencing of this gene results in tissue bleaching, acting as a marker for successful VIGS.

### Gene knock-out using CRISPR/Cas9

To knock-out candidate genes, Cas9-compatible guide RNAs were designed following the parameters previously described (Lin et al., 2023). Two guides were selected for each CRISPR construct. Guide sequences were fused to U6-26 promoters (pICSL90002) and cloned into level-1 vectors (position 3: pICH47751, and position 4: pICH47761). Level-2 constructs were assembled containing the following modules *35S:NPTII:Nos* (pICSL11055, position-1), *AtUbi10:Cas9(with_introns):rbcS-E9t* (pICSL11197, position-2), the guide RNAs (positions −3, −4), and an endlinker (pICH41822). pICSL4723 OD was used as the destination vector.

### Effectoromics screen

To identify effectors recognised by the genes cloned in this study, RXLR effector libraries were screened on *S. americanum* accessions. A library of 311 *P. infestans* effectors was previously screened on these accessions (Lin et al., 2023). Using the predicted proteome of *P. capsici* (Stajich et al., 2021), a new RXLR effector library was generated based on the presence of a signal peptide, RXLR motif, and EER motif. These effectors were cloned into 35S expression vectors (pICSL86977_OD, pICSL86977OD_Aar1, or pICSLUS0004OD), depending on the presence of BsaI, BbsI, or AarI sites. Effectors cloned via the GoldenGate system were fused with a FLAG tag, those cloned through USER cloning remain untagged. *S. americanum* plants were grown under glasshouse conditions, and plants aged four to five week were used for infiltration with *Agrobacterium* suspensions (OD = 0.5). Cell death was assessed at 4 days post-infiltration. For each experiment, two leaves from two different plants were used.

### Plant growth and transformation

*S. americanum* transformation was performed as described by Lin et al. (2023). For each transformation, a minimum of two independent lines are shown. Transgenic *S. lycopersicum* was generated following the previously published protocol (Fillatti et al., 1987), using the cultivar Moneymaker.

*N. benthamiana* was grown in a controlled environment room under conditions of 22°C, 45-65% relative humidity, and a 16-hour light/ 8-hour dark cycle. Both wild type and nrc2/3/4 knockout *N. benthamiana* lines were used (Wu et al., 2019).

### Split luciferase assay

To assess *in planta* association between NLRs and their cognate effectors, a split luciferase assay was performed. NLRs were fused to the C-terminal fragment of luciferase (pICSL50048), while effectors were fused to the N-terminal luciferase fragment (pICSL50047). Coding sequences and tags were cloned into the pICSL86922 vector using GoldenGate assembly. This acceptor contains the CaMV *35S* promoter and *Ocs* terminator. *Agrobacterium* strains carrying these constructs were spot-infiltrated into the *nrc2/3/4 N. benthamiana* line.

Two days post-infiltration, leaves were harvested for imaging. Immediately before imaging, leaves were infiltrated with 100 mM sodium citrate buffer containing 0.4 mM luciferin. Luminescence was visualised using the Nightowl II LB 983 In vivo imaging system and Winlight software (Berthold Technologies, Germany). As a positive control, the *S. americanum* NLR Rpi-amr3 and its cognate effector Avramr3 were used (Witek et al., 2016, Lin et al., 2022).

### Disease assays

Two *P. infestans* strains were used: strain 88069 for infection of *S. americanum* and strain 90128 for infection of *S. lycopersicum*. Both strains were maintained at 17 °C on rye-sucrose agar (RSA) medium. To induce zoospore release, 14-day-old plates were flooded with cold water and incubated at 4 °C for 1-2 hours. Zoospore concentrations were determined using a haemocytometer before dilution to the appropriate concentration (as indicated in figure legends).

For *P. capsici* infection assays, LT1534 was used. *P. capsici* was maintained on V8 agar plates incubated at 25 °C. To induce zoospore release, 10-day-old plates were flooded with cold water and incubated at 4 °C for 1 hour, followed by 1 hour at room temperature. The suspension was diluted to 10,000 spores ml^-1^.

Detached leaf assays were used to assess the resistance of *S. americanum* and *S. lycopersicum* to *P. infestans* and *P. capsici*. Fully expanded leaves from five- to six-week-old plants were placed on damp paper within 500 cm^2^ square culture dishes. Leaves were inoculated by either transfer of infected agar plugs (2-3 mm^2^ plugs were used in quantitative *P. capsici* infection assays), or by pipetting 10 µl droplets onto each leaf. For *P. infestans* infection assays, trays were incubated at 17 °C for 5-10 days until susceptible controls showed infection. For *P. capsici* assays, trays were incubated at 25 °C for 2-4 days until susceptible controls showed infection.

### BSA-RenSeq and map-based cloning

Three F2 mapping populations were used in this study: (1) SP2271 x SP2275, (2) SP2271 x SP2308, and (3) SP2271 x SP2297. Population 1 was phenotyped for *P. infestans* resistance, population 2 for *P. capsici* resistance, and population 3 for responsiveness to transient expression of PITG_16275. Genomic DNA was extracted from leaf tissue using the Qiagen DNeasy plant kit (Qiagen, 69104). RenSeq libraries were then prepared using bulked genomic DNA from resistant and susceptible individuals. Enrichment for NLR-encoding genes was performed using the V4 RenSeq bait library (Witek et al., 2021). Libraries were sequenced using paired 250 bp illumina sequencing by Novogene (Beijing, China). Bulked segregant analysis was performed following previously described methods (Jupe et al., 2013, Andolfo et al., 2014, Witek et al., 2016). Resistance-linked *R*-gene candidates were filtered based on the presence of motifs identified using NLR-parser, and, when available, by their expression as determined using cDNA RenSeq. To detect expression, cDNA RenSeq reads were mapped using HISAT spliced aligner under default settings (Kim et al., 2015). BAM files were visualised using IGV to assess expression and detect splice variants.

### Annotation of NLRs and gene models

The NLR genes either within mapping intervals, or on resistance-linked RenSeq contigs were predicted using NLR-parser (Steuernagel et al., 2015). To generate accurate gene models, cDNA RenSeq reads were mapped to reference sequences as described above. Resulting BAM files were manually inspected to annotate NLR-encoding genes. Published sequences and gene models for *S. americanum* genomes were used to support the manual annotation.

## Author contributions

R.H., K.W., X.L., and J.D.G.J conceived and designed the project. R.H., A.C.O.A., M.S., L.L., H.K., M.A., S.P., V.A., K.W., and X.L. performed the experiments. R.H., K.W., and X.L. performed the bioinformatic analyses. R.H., L.L., and X.L. contributed to the cloning of the *P. capsici* effector library, and the effector screening in *S. americanum.* M. Smoker, J.T., A.W-K., and K.B. performed plant transformation and tissue culture. R.H., X.L., and J.D.G.J wrote the manuscript with input from all authors. K.H.S., K.W., X.L., and J.D.G.J. contributed resources. All authors approved the manuscript.

## Competing interests

The authors declare no competing interests.

## Acknowledgements

This research was financed by BBSRC grants BB/P021646/1 (J.D.G.J.), BB/S018832/1 (J.D.G.J.), BB/W017423/1 (J.D.G.J.), the Gatsby Charitable Foundation (J.D.G.J), and the National Natural Science Foundation of China (32441060 and 32372506, X.L.). We thank the TSL transformation and tissue culture team, the SynBio team, media services, bioinformatics team, and the horticulturists (Sara Perkins, Justine Smith, Catherine Taylor, Timothy Wells, and Matt Castle) who supported this work. We also thank Adeline Harrant and Sophien Kamoun for supplying the *P. infestans* strain 90128.

## Supplementary Material

**Supplementary figure 1.** *Rpi-amr5* candidates were named by their homology to members of the SP2273 *Rpi-amr1* cluster.

**Supplementary figure 2.** Resistance in SP2275 is not due to *Rpi-amr1*.

**Supplementary figure 3.** VIGS of the *Rpi-amr1c* paralog compromises *P. infestans*resistance in SP2275.

**Supplementary figure 4.** Transient expression of *Rpi-amr5* and *Rpc2* does not result in constitutive activity in *S. americanum*.

**Supplementary figure 5.** CRISPR/Cas9-mediated mutation of *Rpi-amr5* in *S. americanum*.

**Supplementary figure 6.** *Rpc2* candidates were named by their homology to members of the SP2273 *Rpi-amr1* cluster.

**Supplementary figure 7.** VIGS of the *Rpi-amr1c* paralog compromises *P. capsici* resistance in SP2308.

**Supplementary figure 8.** CRISPR-induced knockout of *Rpi-amr5* does not elevate susceptibility to *P. capsici*.

**Supplementary figure 9.** Constitutive activity of Rpi-amr5 and Rpc2 in *N. benthamiana* is observed following expression under an NLR promoter.

**Supplementary figure 10.** CRISPR/Cas9-mediated mutation of *Rpc2* in *S. americanum*.

**Supplementary figure 11.** Expanded Figure 4b.

**Supplementary figure 12.** AVRamr1 amino acid alignment.

**Supplementary figure 13.** Alignment of Rpi-amr1-2273 and Rpi-amr1-2308 amino acid sequences.

**Supplementary figure 14.** Rpi-amr5 and Rpc2 are constitutively active when expressed with NbNRC2/NbNRC3/SaNRC3, but not SaNRC2.

**Supplemental figure 15.** *Rpi-amr5* function has been validated in three *S. americanum* accessions.

**Supplementary figure 16.** Rpi-amr5 and Rpc2 can signal through either SaNRC1, or SaNRC2.

**Supplemental figure 17.** Structural homologs of PITG_16275 are found in several *Phytophthora* species, but not *P. capsici*.

**Supplementary figure 18.** Rpi-amr5 recognises PITG_16275 orthologs from *P. parasitica*, *P. megakarya* and *P. palmivora*.

**Supplementary figure 19.** Rpi-amr5 and Rpc2 recognise three *P. capsici* effectors with low sequence similarity to PITG_16275 or PITG_09160.

**Supplementary figure 20.** PITG_09160 recognition is reported in several *S. americanum* accessions that lack complete *Rpi-amr5* alleles.

**Supplementary table 1.** Four NLR encoding genes were identified as *Rpi-amr5* candidates.

**Supplementary table 2.** Five NLR encoding genes were identified as *Rpc2* candidates.

**Supplementary table 3.** Effectors recognised by Rpi-amr1e-2308, Rpi-amr5, or Rpc2.

**Supplementary table 4.** NLR-encoding genes characterized in this study.

## Notes

### Competing Interest Statement

The authors have declared no competing interest.

