## Supplementary figures and tables for "Diverse haplotypes at a complex *Solanum americanum* locus confer resistance to *Phytophthora infestans* and *P. capsici*"

* Co-corresponding author

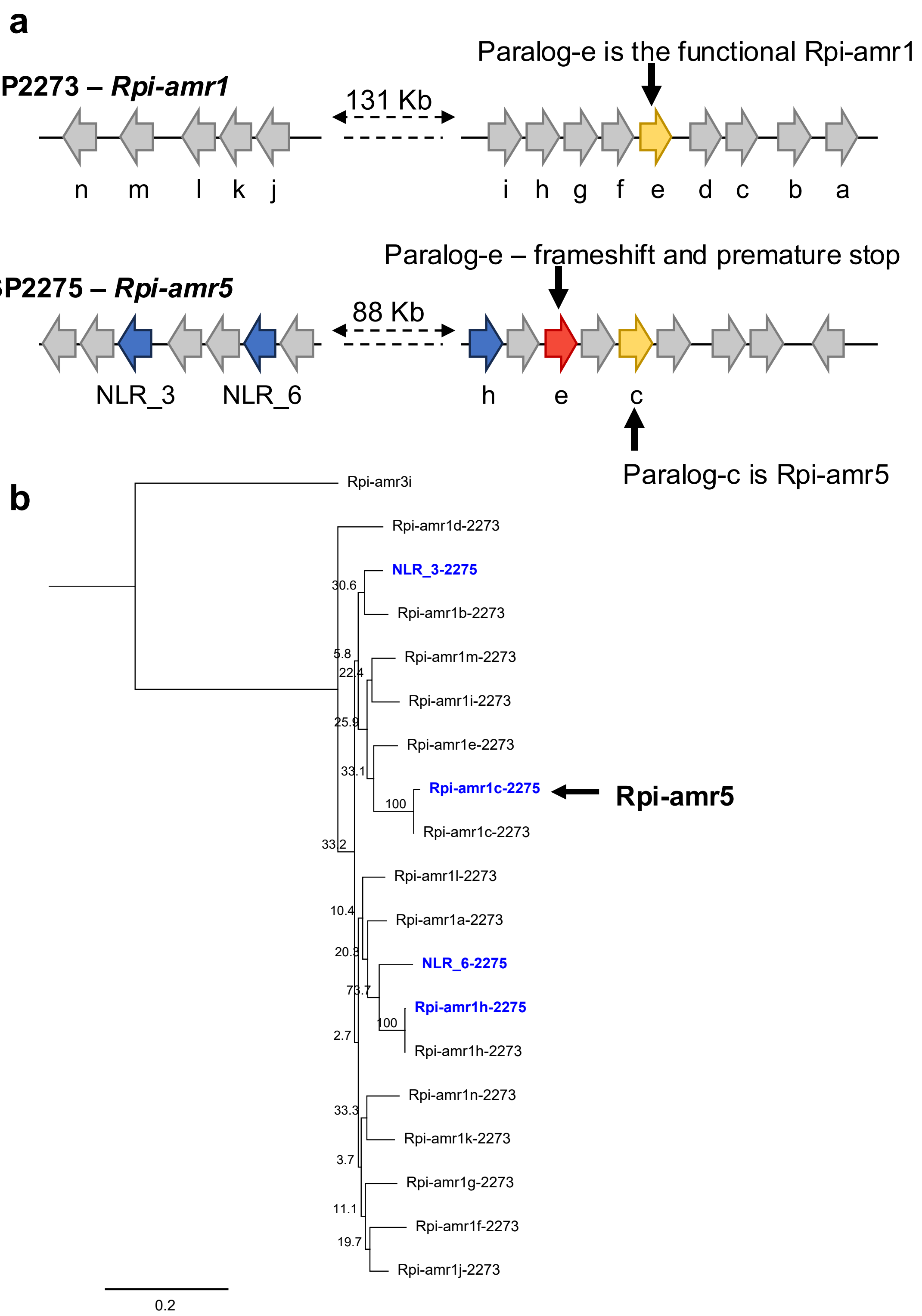


**Supplementary figure 1. *Rpi-amr5* candidates were named by their homology to members of the SP2273 *Rpi-amr1* cluster. (a)** The *Rpi-amr1* clusters in *S. americanum* accessions SP2273 (the original source of *Rpi-amr1*) and SP2275 (the source of *Rpi-amr5*). NLR-encoding genes are represented using arrows, genes which confer *P. infestans* resistance are coloured yellow, other *Rpi-amr5* candidates are coloured blue. The *Rpi-amr1* ortholog (*Rpi-amr1e*, indicated in red) is non-functional in SP2275 due to a premature stop codon. **(b)** Phylogenetic tree showing the relationship of the Rpi-amr5 candidates with NLRs of the Rpi-amr1 cluster from the accession SP2273. This tree was used to name two candidates which have close homology to SP2273 NLRs, these are Rpi-amr1c-2275 and Rpi-amr1h-2275. The other two candidates were generically named NLR_3-2275 and NLR_6-2275 based on their position within the SP2275 cluster. The functional Rpi-amr5 is indicated. The tree is rooted using Rpi-amr3.

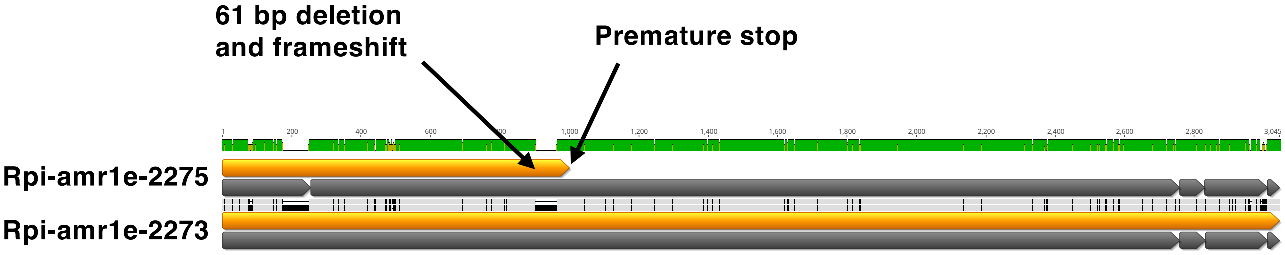


**Supplementary figure 2. Resistance in SP2275 is not due to *Rpi-amr1*.** Resistance in SP2275 is not due to *Rpi-amr1*. The *Rpi-amr1e* paralog, which is responsible for resistance in SP2273, is non-functional in SP2275 due to a frameshift and premature stop. The position of this stop is indicated in the alignment. The CDS of each allele is shown, grey bars indicate the position of exons and orange bars the ORF.

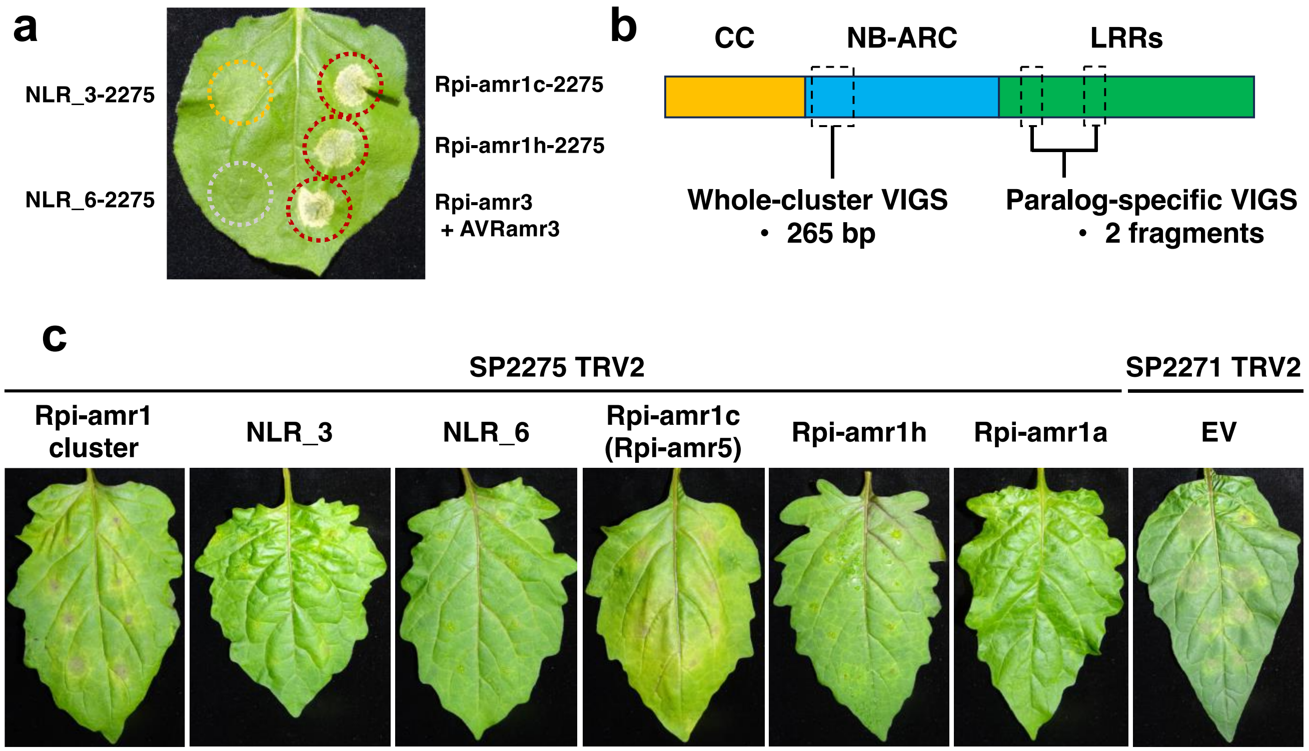


**Supplementary figure 3. VIGS of the *Rpi-amr1c* paralog compromises *P. infestans* resistance in SP2275. (a)** Transient expression of *Rpi-amr1c-2275* and *Rpi-amr1h-2275* in *N. benthamiana* results in strong HR indicating constitutive activity. *N. benthamiana* infiltrations were performed using *Agrobacterium* suspensions at 0.2 $OD_{600}$, leaves were imaged at 4 dpi. **(b)** VIGS was used to test the functionality of *Rpi-amr5* candidates. Whole-cluster silencing was performed using a fragment from the NB-ARC encoding region of the *Rpi-amr1h* paralog. Paralog-specific silencing was performed using constructs containing two LRR-encoding fragments fused together. **(c)** Silencing of the *Rpi-amr1* cluster and the *Rpi-amr1c* paralog compromised resistance in SP2275, silencing of *Rpi-amr1a* or *Rpi-amr1h* did not affect resistance. Leaves were inoculated with 10 µl drops of *P. infestans* (strain 88069) spores at a concentration of 20,000 spores $mL^{-1}$. Leaves were photographed at 5 days post-inoculation.

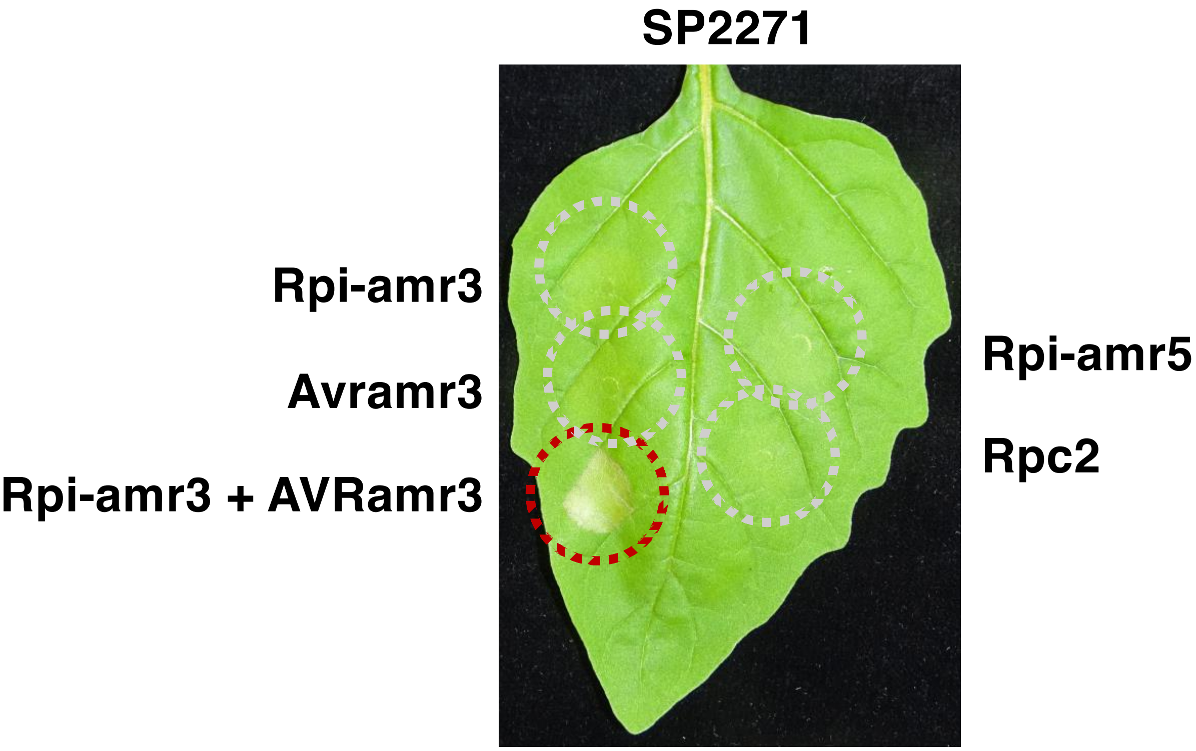


**Supplementary figure 4. Transient expression of *Rpi-amr5* and *Rpc2* does not result in constitutive activity in *S. americanum*.** *Agrobacterium*-mediated transient expression of *Rpi-amr5* and *Rpc2* does not result in HR in *S. americanum* accession SP2271. *S. americanum* infiltrations were performed at 0.5 $OD_{600}$, leaves were imaged at 3 dpi.

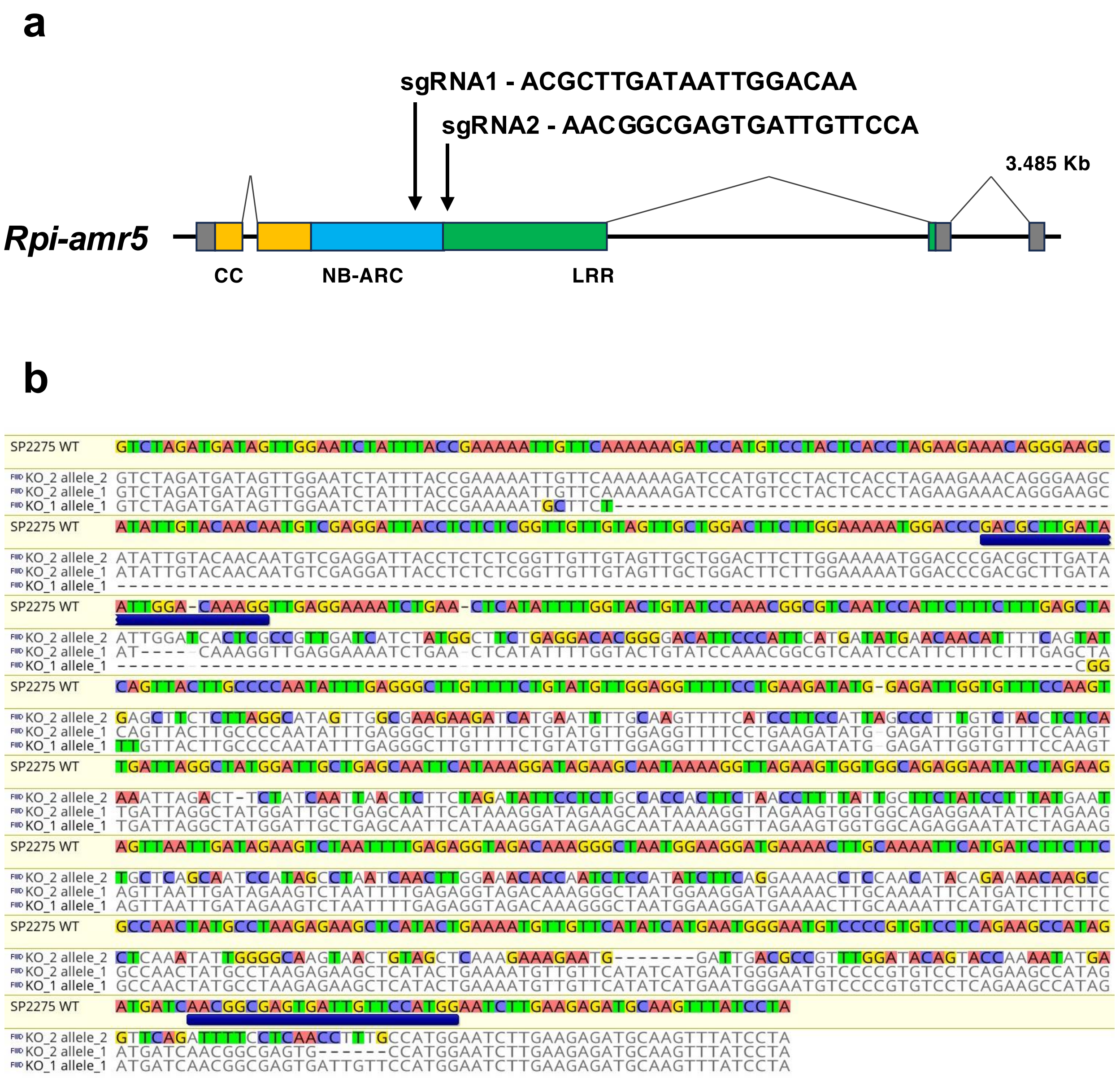


**KO_1 allele_1 – 205 bp deletion at first sgRNA site**

**KO_2 allele_1 – Indels at both sgRNA sites (-4, -6)**

**KO_2 allele_2 – Inversion between sgRNA sites**

**Supplementary figure 5. CRISPR/Cas9-mediated mutation of *Rpi-amr5* in *S. americanum*. (a)** To confirm the function of *Rpi-amr5* in SP2275, knockout lines were produced using CRISPR/Cas9. Two sgRNAs were designed to target *Rpi-amr5*, their positions within the gene are indicated by black arrows. **(b)** Two independent *Rpi-amr5* knockout lines that exhibited a loss of *P. infestans* resistance were selected for further analysis. To genotype these mutants, a short genomic region spanning the CRISPR target sites was amplified and cloned. For each line, six independent colonies were purified and sequenced. The resulting genotypes are shown, with polymorphisms relative to the SP2275 reference highlighted. The positions of sgRNAs within this region are indicated by blue boxes.

a

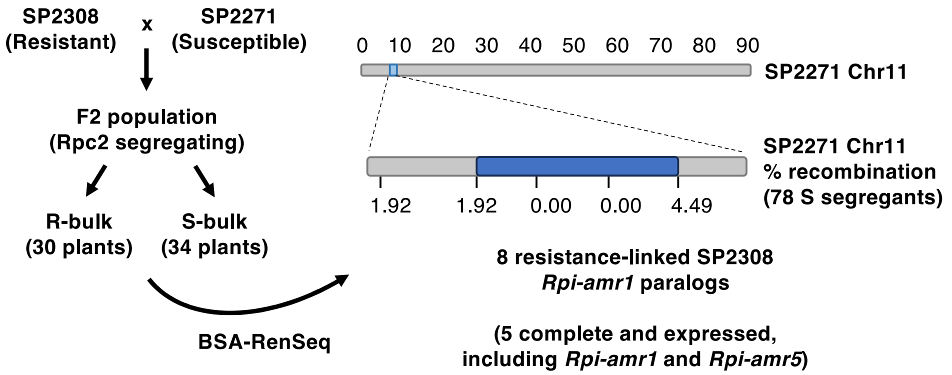


b

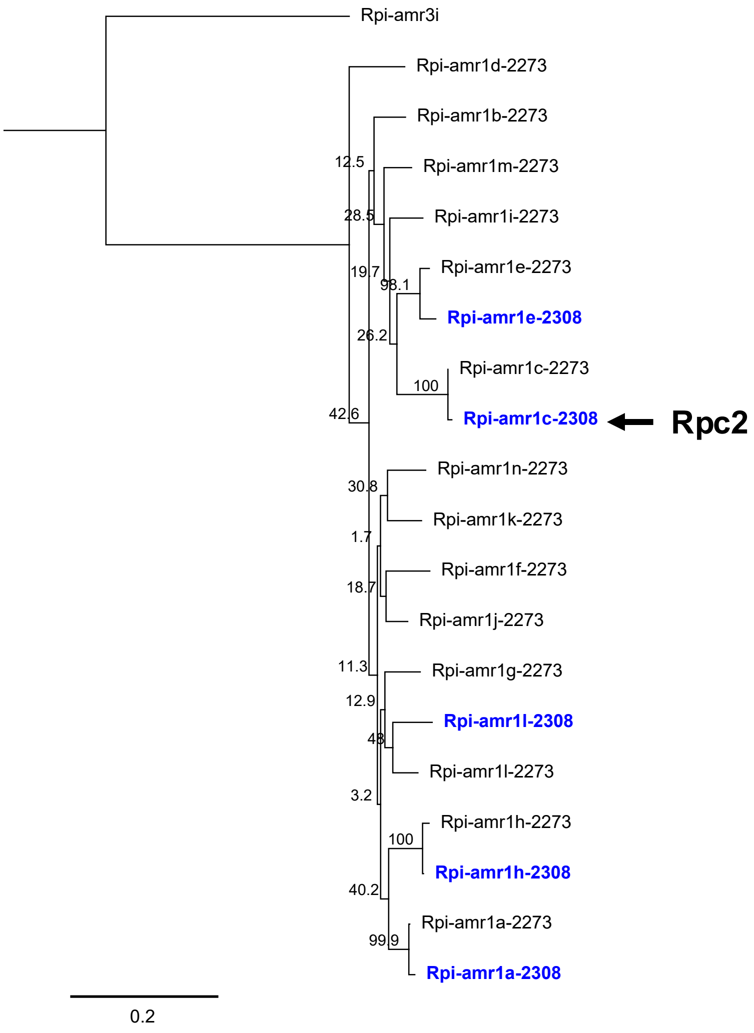


**Supplementary figure 6. *Rpc2* candidates were named by their homology to members of the SP2273 *Rpi-amr1* cluster. (a)** SP2308 was crossed to SP2271 and an F2 population was phenotyped, the 34 most susceptible and 30 most resistant plants were used for bulked segregant analysis. Using the SP2271 reference genome, *Rpc2* was mapped to the *Rpi-amr1* cluster on Chromosome 11. Fine mapping in an expanded population of 78 susceptible segregants was performed, percentage recombination rates are indicated. Eight NLR-encoding genes within this interval were identified in the SP2308 SMRT RenSeq assembly, the five complete candidates are indicated in blue. *Rpi-amr1* and *Rpi-amr5* are both complete in SP2308. **(b)** Phylogenetic tree showing the relationship between Rpc2 candidates and NLRs from the *Rpi-amr1* cluster in *S. americanum* accession SP2273. This tree was used to name the candidates – Rpi-amr1a-2308, Rpi-amr1c-2308, Rpi-amr1e-2308, Rpi-amr1h-2308 and Rpi-amr1l-2308. The functional Rpc2 is indicated. The tree is rooted using Rpi-amr3.

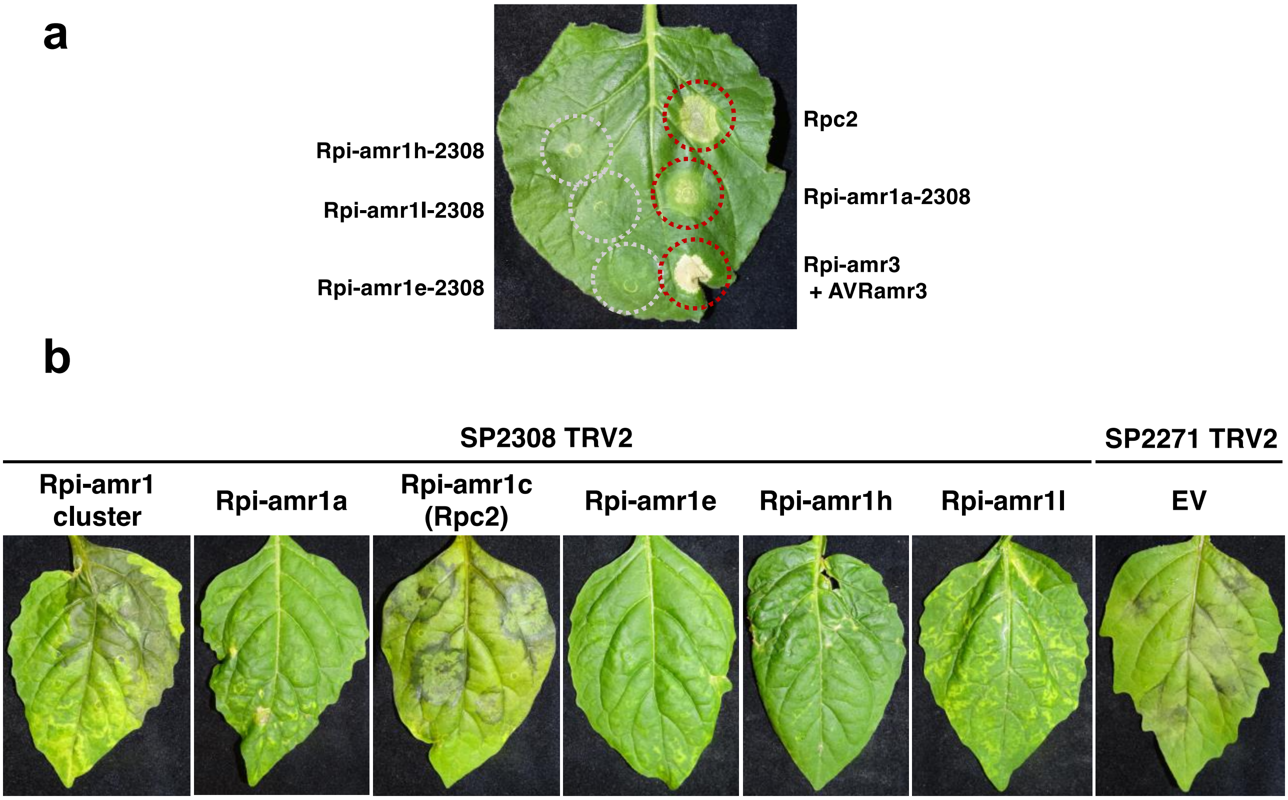


**Supplementary figure 7. VIGS of the *Rpi-amr1c* paralog compromises *P. capsici* resistance in SP2308. (a)** *Rpc2* and the four complete paralogs within the SP2308 *Rpi-amr1* cluster were cloned into CaMV *35S* expression vectors to be tested in *N. benthamiana*. Transient expression of both *Rpi-amr1c-2308* and *Rpi-amr1a-2308* produced strong HR indicating constitutive activity. *N. benthamiana* infiltrations were performed at 0.2 $OD_{600}$, leaves were imaged at 4 dpi. **(b)** VIGS was performed to silence candidates in SP2308. Silencing of either the whole cluster, or the *Rpi-amr5* paralog (*Rpi-amr1c*) compromised resistance in SP2308. Silencing of *Rpi-amr1e-2308* did not result in susceptibility. Leaves were inoculated with 10 µl drops of *P. capsici* strain LT1534 at a concentration of 20,000 spores $ml^{-1}$ and imaged at 3 days post-inoculation.

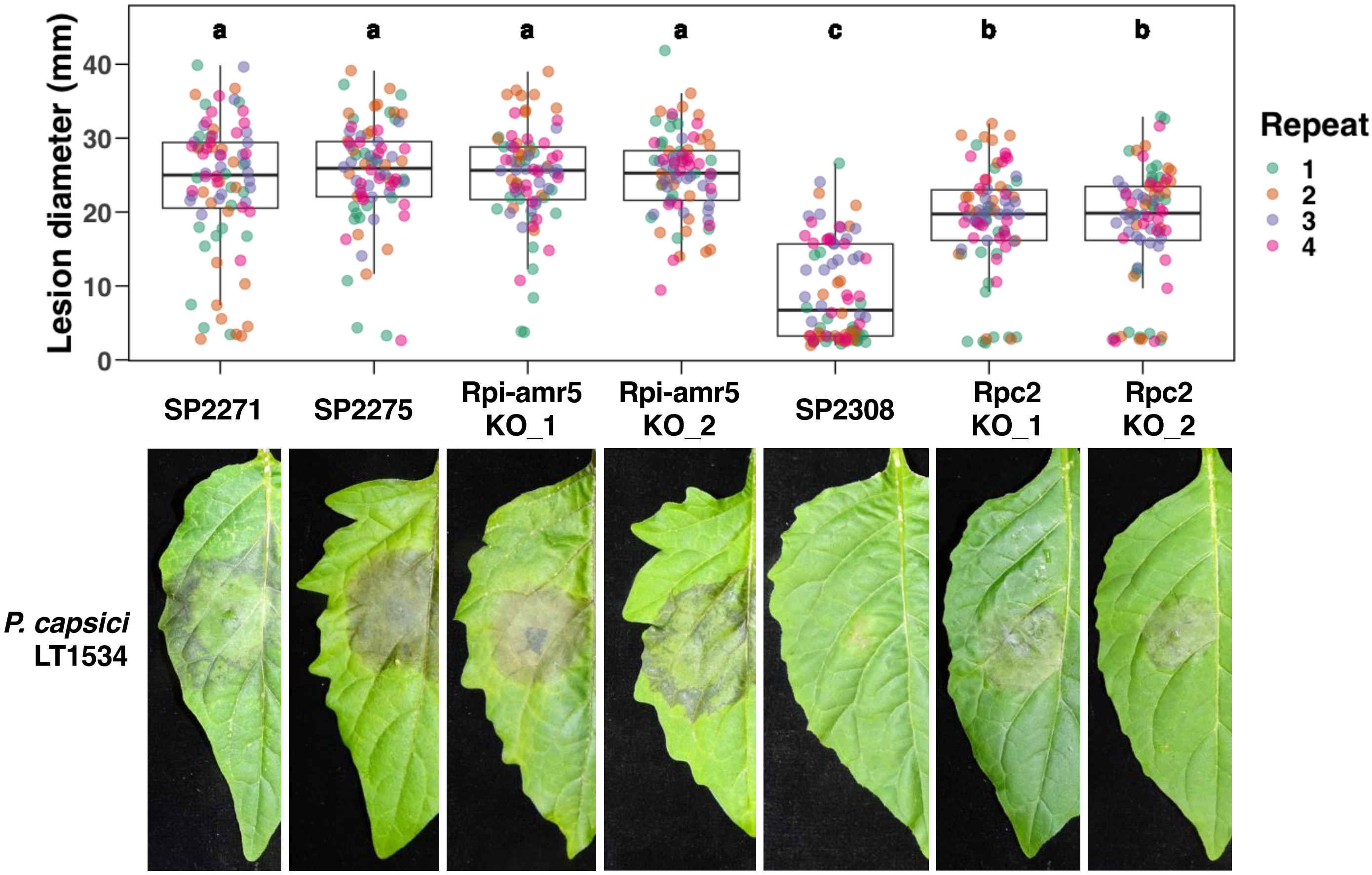


**Supplementary figure 8. CRISPR-induced knockout of *Rpi-amr5* does not elevate susceptibility to *P. capsici*.** CRISPR-induced knock out of *Rpi-amr5 (Rpi-amr1c)* in SP2275 does not further increase susceptibility to *P. capsici*. Resistance in all SP2275 lines is indistinguishable from susceptible accession SP2271. Leaves were inoculated by transfer of *P. capsici* (strain LT1534) mycelial plugs and imaged 3 days post-inoculation. Two independent biallelic mutants are shown. Four biological replicates were performed; all data points (80 per treatment) are represented as box-and-whisker plots. Statistical differences determined using one-way ANOVA and Tukey's HSD test ($p < 0.05$).

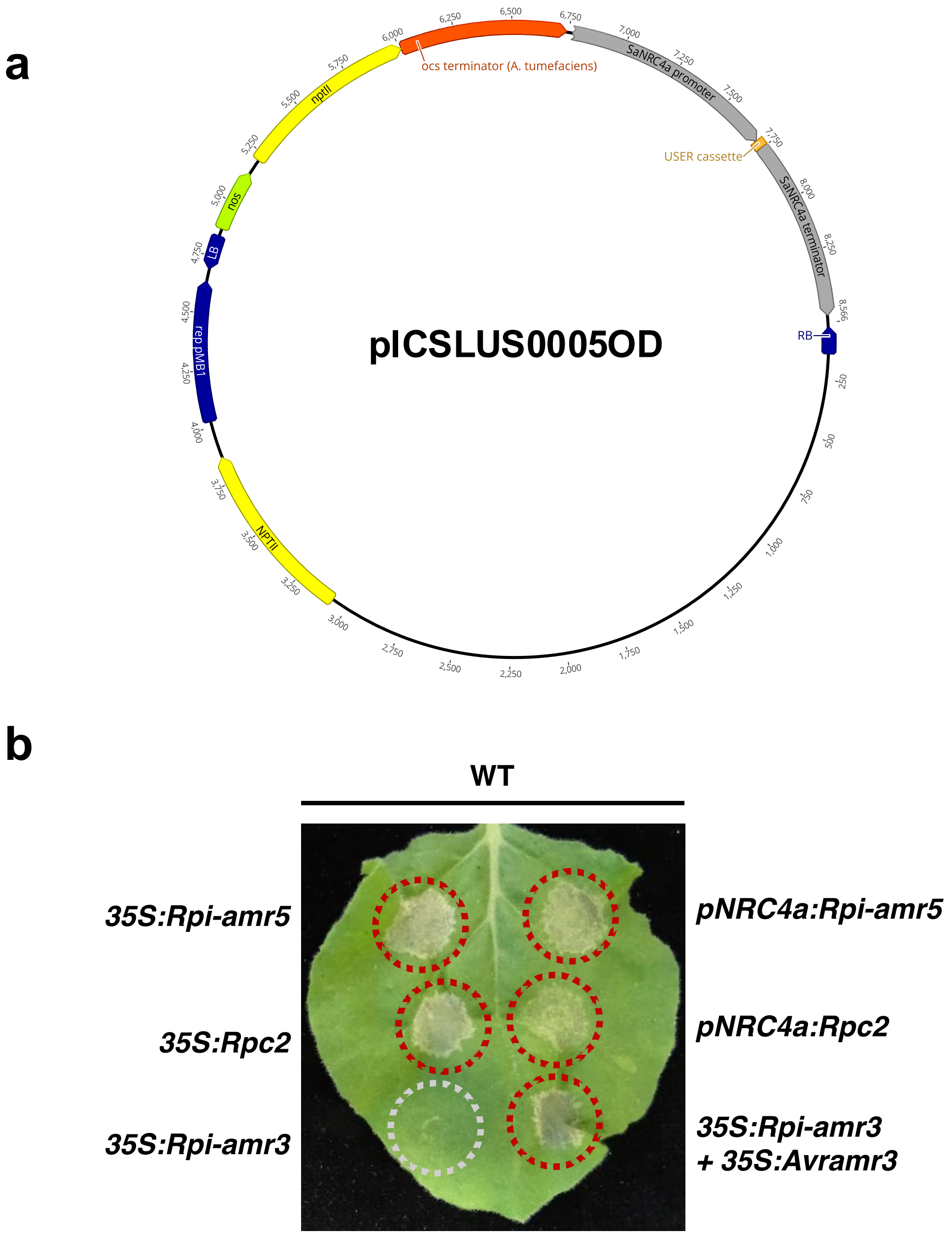


**Supplementary figure 9. Constitutive activity of Rpi-amr5 and Rpc2 in *N. benthamiana* is observed following expression under an NLR promoter. (a)** Custom USER vector pICSLUS0005OD was designed to express NLRs under a moderate promoter. The *SaNRC4a* promoter (943 bp) and terminator (793 bp) was cloned from the *S. americanum* reference accession SP1102. A USER cassette was fused between these sequences to facilitate linearisation of the plasmid and insertion of target sequences. **(b)** *Rpi-amr5* and *Rpc2* confer constitutive activity in *N. benthamiana* under both CaMV *35S* and *S. americanum NRC4a* regulatory elements. Clear cell death was visible after 3 days. Cell death is equivalent to effector-dependent activation of Rpi-amr3. *Agrobacterium* suspensions were infiltrated at 0.2 $OD_{600}$.

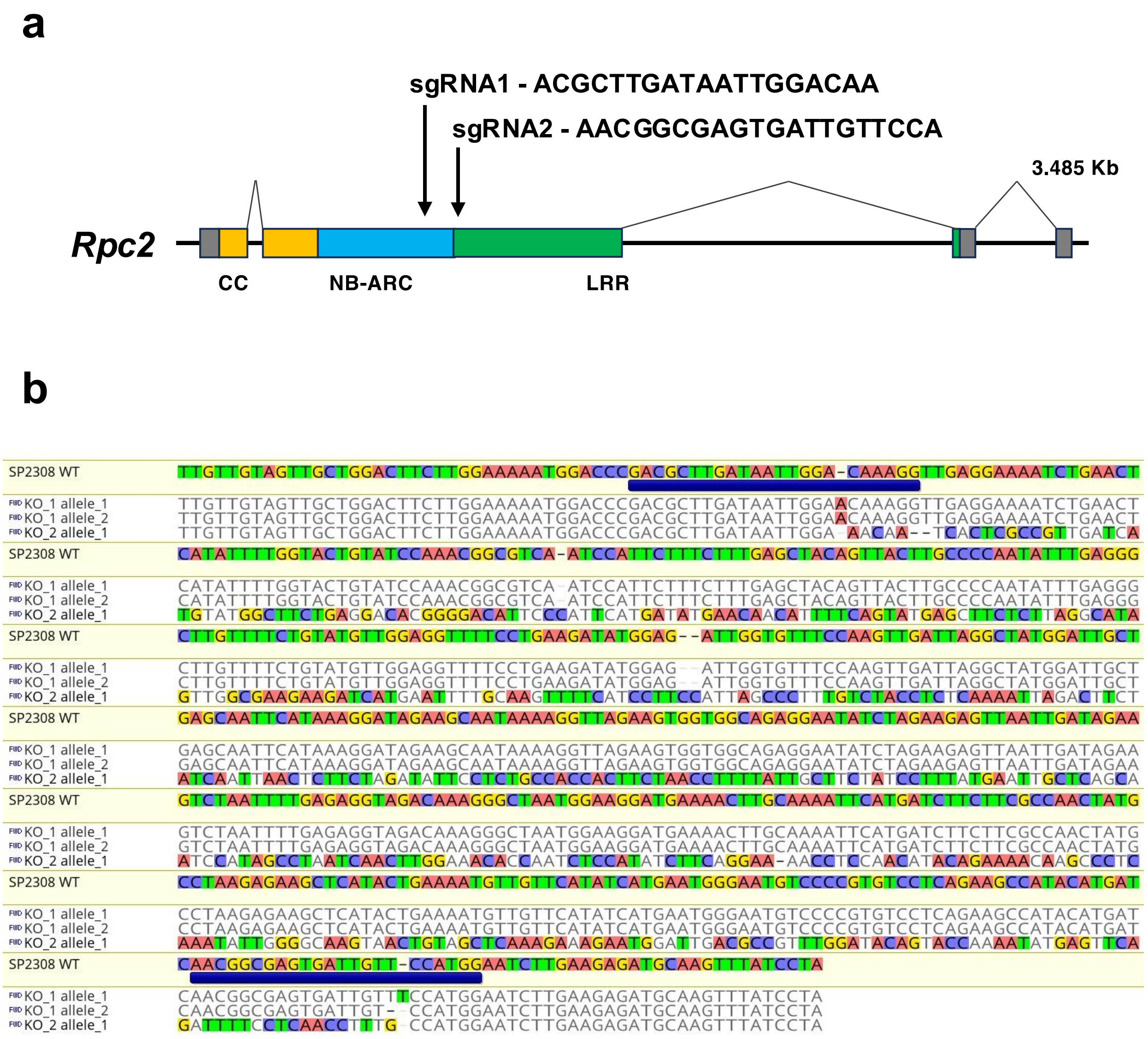


**KO_1 allele_1 – Indels at both sgRNA sites (+1, +1)**

**KO_1 allele_2 – Indels at both sgRNA sites (+1, -1)**

**KO_2 allele_1 – Inversion between sgRNA sites**

**Supplementary figure 10. CRISPR/Cas9-mediated mutation of *Rpc2* in *S. americanum*.** **(a)** To confirm the function of *Rpc2* in SP2275, knockout lines were produced using CRISPR/Cas9. Two sgRNAs were designed to target *Rpc2*, their positions within the gene are indicated by black arrows. **(b)** Two independent *Rpc2* knockout lines that exhibited a loss of *P. capsici* resistance were selected for further analysis. To genotype these mutants, a short genomic region spanning the CRISPR target sites was amplified and cloned. For each line, six independent colonies were purified and sequenced. The resulting genotypes are shown, with polymorphisms relative to the SP2308 reference highlighted. The positions of sgRNAs within this region are indicated by blue boxes.

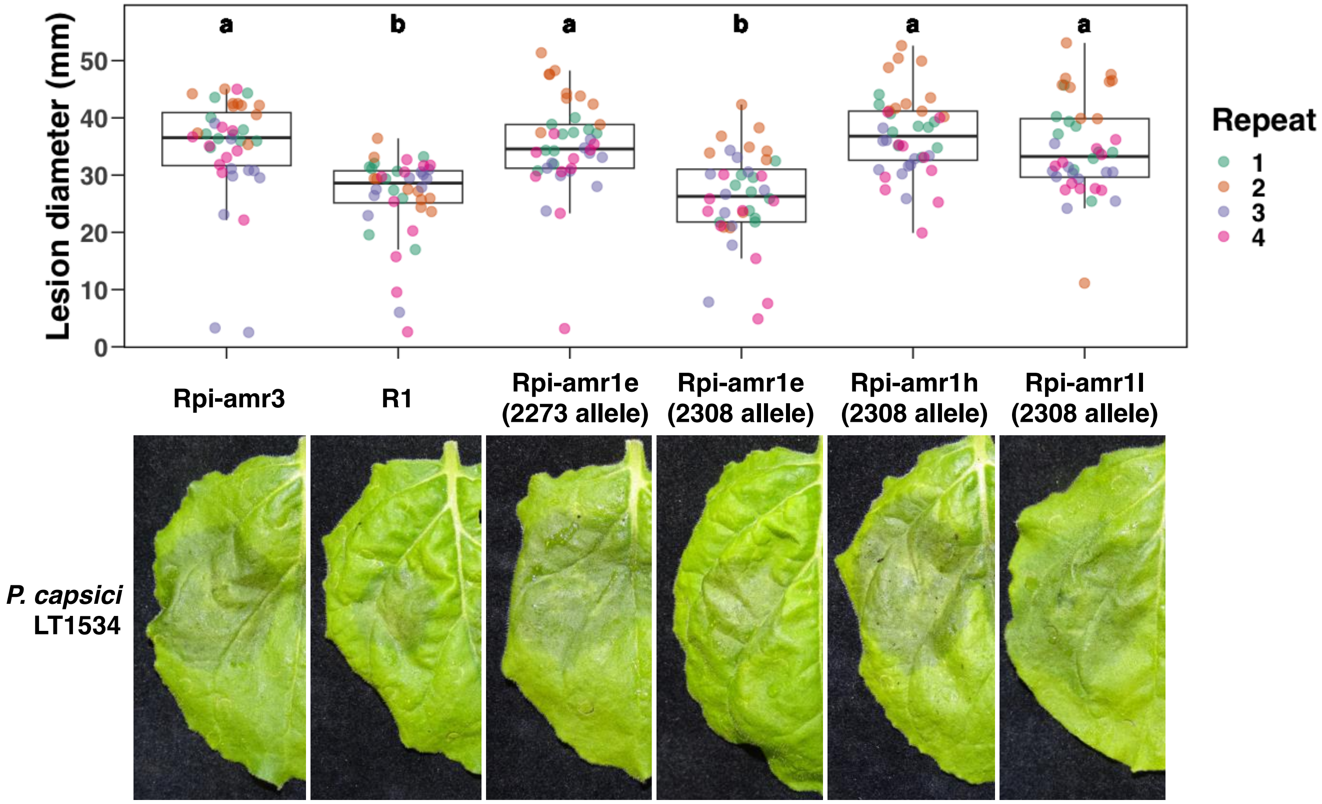


**Supplementary figure 11. Expanded Figure 4b.** *Rpi-amr1e-2308* confers partial resistance to *P. capsici* in *N. benthamiana* transient assays. Other non-constitutively active paralogs from the *Rpi-amr1* cluster in SP2308 do not. *R1* was used as a positive control for *P. capsici* resistance. Agroinfiltration was performed using suspensions normalised to 0.5 $OD_{600}$. At 2 dpi, leaves were detached and *P. capsici* mycelial plugs were transferred onto the leaves. Lesion size was measured at 3 days post-inoculation. Four biological replicates were performed, all data points (40 per treatment) are represented as box and whisker plots. Statistical differences determined using one-way ANOVA and Tukey's HSD test ($p < 0.05$).

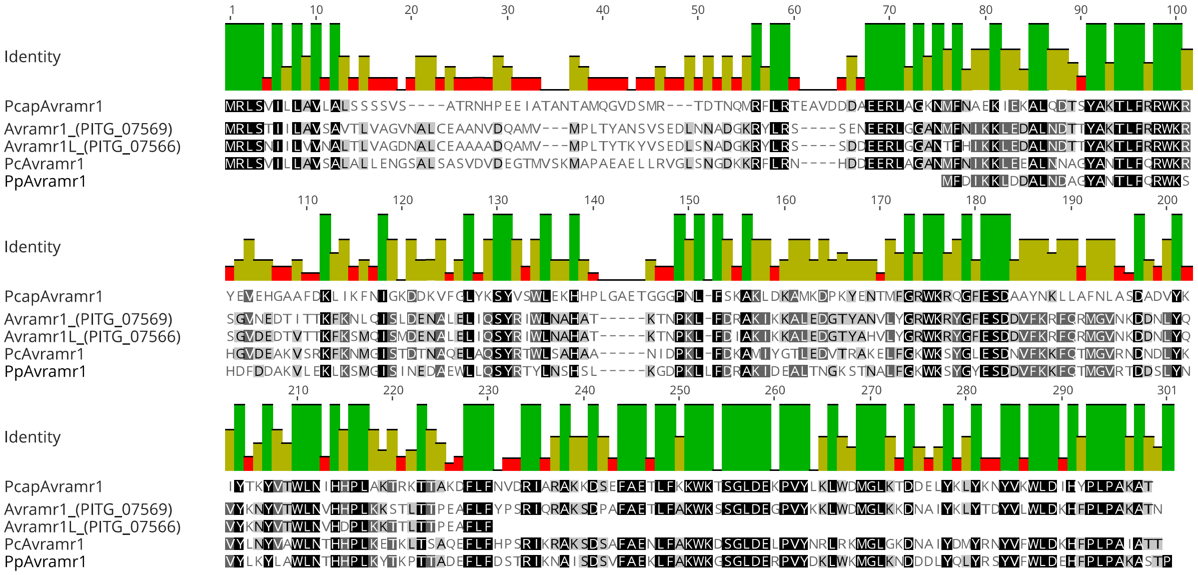


**Supplementary figure 12. AVRamr1 amino acid alignment.** The alignment of AVRamr1 protein sequences from *Phytophthora infestans* (Pi), *Phytophthora cactorum* (Pc), *Phytophthora parasitica* (Pp) and *Phytophthora capsici* (Pcap). PcapAVRamr1 was identified in this study, other alleles were identified by Lin et al. 2020.

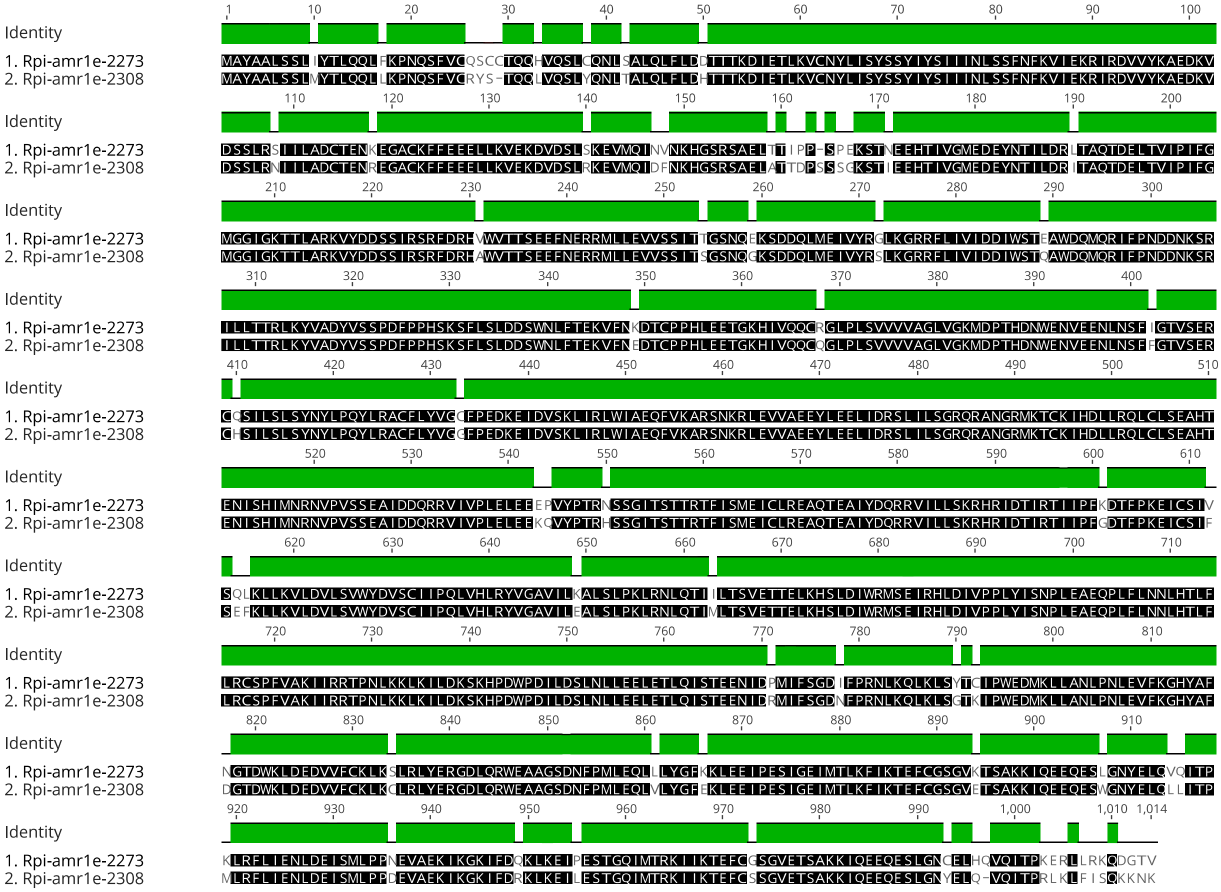


**Supplementary figure 13. Alignment of Rpi-amr1-2273 and Rpi-amr1-2308 amino acid sequences.** The protein alignment of Rpi-amr1 sequences from *S. americanum* accessions SP2273 and SP2308. The SP2273 allele was characterised by Witek et al 2021, the SP2308 allele was cloned in this study. The *Rpi-amr1-2308* gene model was predicted through homology to the published *Rpi-amr1-2273* model.

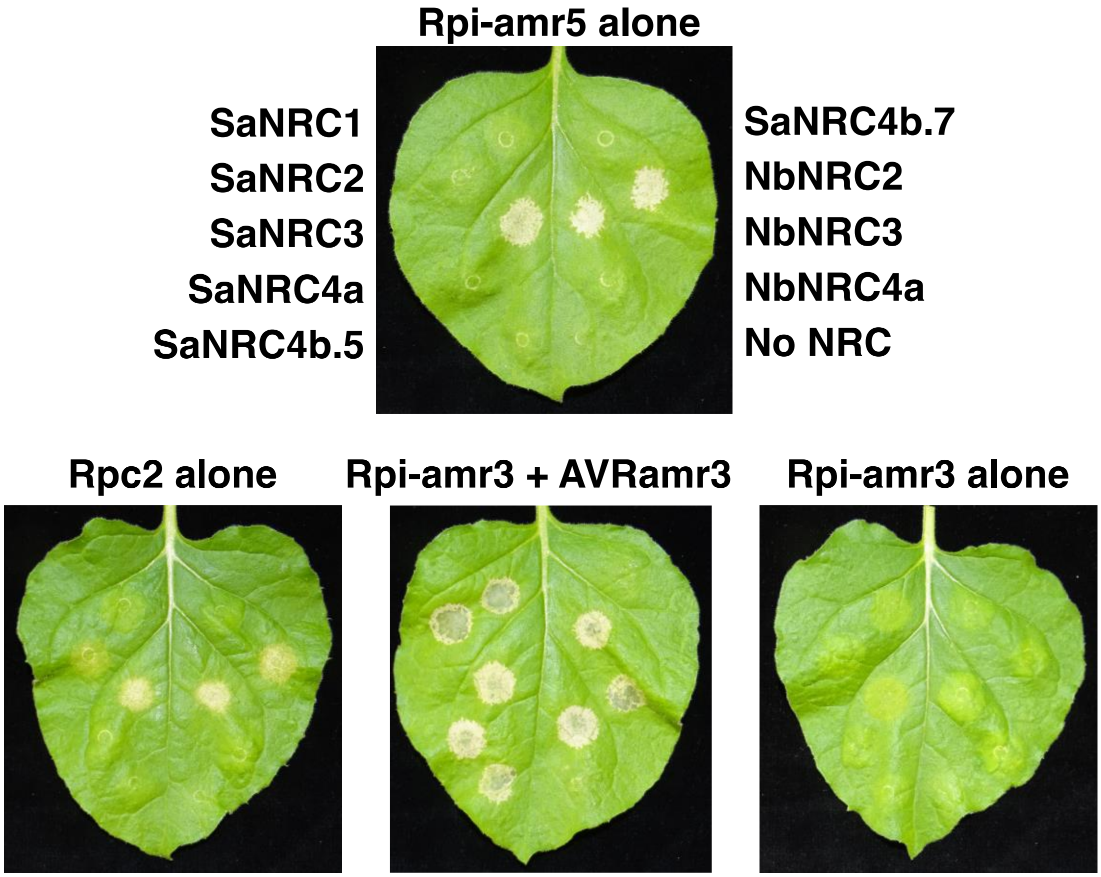


**Supplementary figure 14. Rpi-amr5 and Rpc2 are constitutively active when expressed with NbNRC2/NbNRC3/SaNRC3, but not SaNRC2.** Absence of NRC helper NLRs abolishes Rpi-amr5/Rpc2 constitutive activity. Complementation with NbNRC2, NbNRC3, or SaNRC3 restores constitutive activity. Transient expression was performed in the nrc2/3/4 knock out *N. benthamiana* line. *Agrobacterium* strains were infiltrated at 0.2 $OD_{600}$ and leaves were imaged at 3 dpi.

**a**

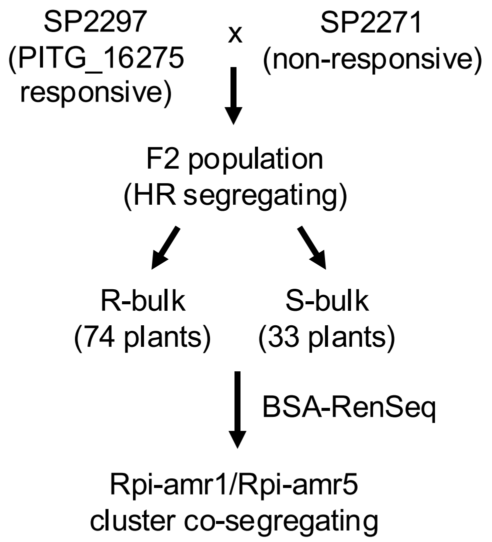


**b**

| | Rpi-amr1e-... | Rpc-amr1c-... | Rpi-amr1c-... | Rpi-amr1c-... |
|---|---|---|---|---|
| Rpi-amr1e-2273 | | 74.0% | 74.1% | 74.1% |
| Rpc-amr1c-2308 | 74.0% | | 97.4% | 97.4% |
| Rpi-amr1c-2275 | 74.1% | 97.4% | | 100% |
| Rpi-amr1c-2297 | 74.1% | 97.4% | 100% | |

**Supplemental figure 15. *Rpi-amr5* function has been validated in three *S. americanum* accessions. (a)** Recognition of PITG_16275 was mapped in an F2 population produced by crossing the responsive line SP2297 with the non-responsive line SP2271. Responsiveness segregates in a 3:1 ratio of responsive: non-responsive (121 R and 33 NR; $\chi 2$ (1, N = 154) = 0.39 P = 0.3061). Bulked segregant analysis was used to map the recognition to the *Rpi-amr1*/*Rpi-amr5*/*Rpc2* cluster on Chromosome 11. **(b)** Functionally validated *Rpi-amr1c* alleles encode NLRs with high amino acid identity. Heatmap of the percent identity of aligned amino acid sequence of these alleles. All three were independently mapped using either resistance to *P. infestans* (*Rpi-amr1c-2275*), resistance to *P. capsici* (*Rpi-amr1c-2308*), or recognition of PITG_16275 (*Rpi-amr1c-2297*).

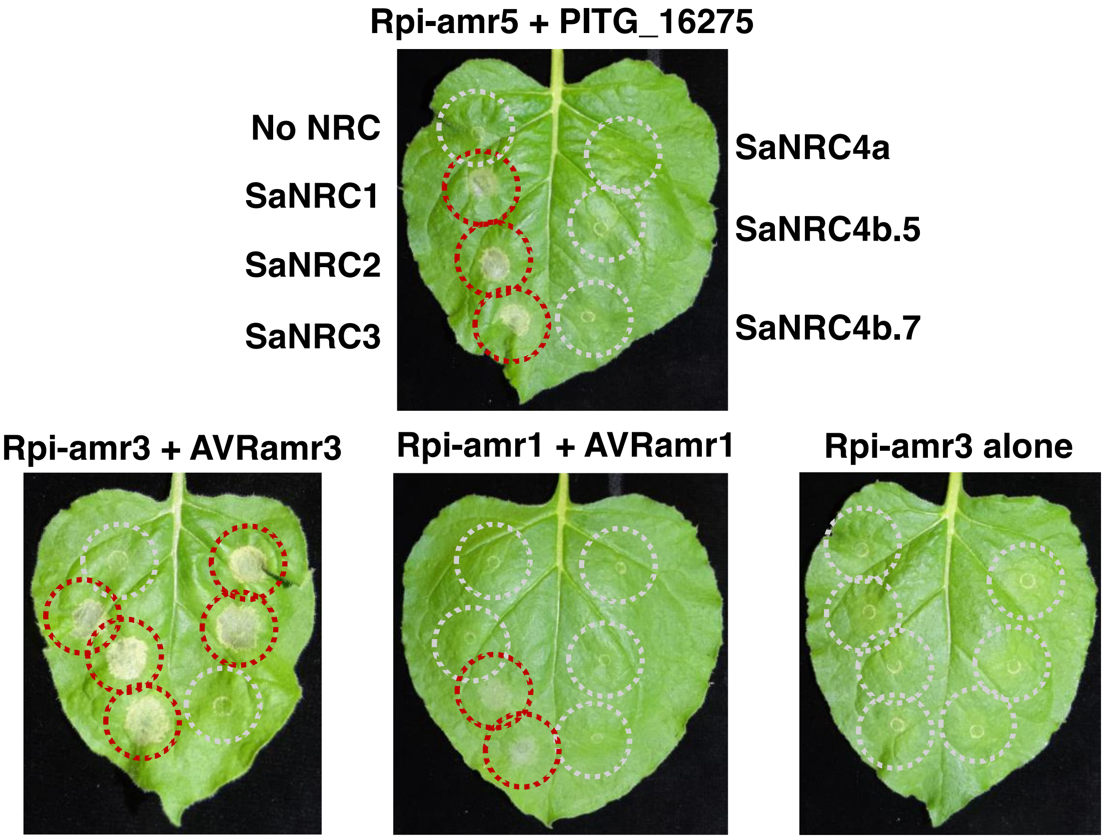


**Supplementary figure 16. Rpi-amr5 and Rpc2 can signal through either SaNRC1, or SaNRC2.** PITG_16275-mediated activation of Rpi-amr5 and Rpc2 triggers SaNRC1- and SaNRC2- dependent cell death. SaNRC3 mediated cell death cannot be distinguished from effector-independent constitutive activity (demonstrated in Fig. S14). Transient expression was performed in the nrc2/3/4 knock out *N. benthamiana* line. *Agrobacterium* strains were infiltrated at 0.2 $OD_{600}$ and leaves were imaged at 3 dpi.

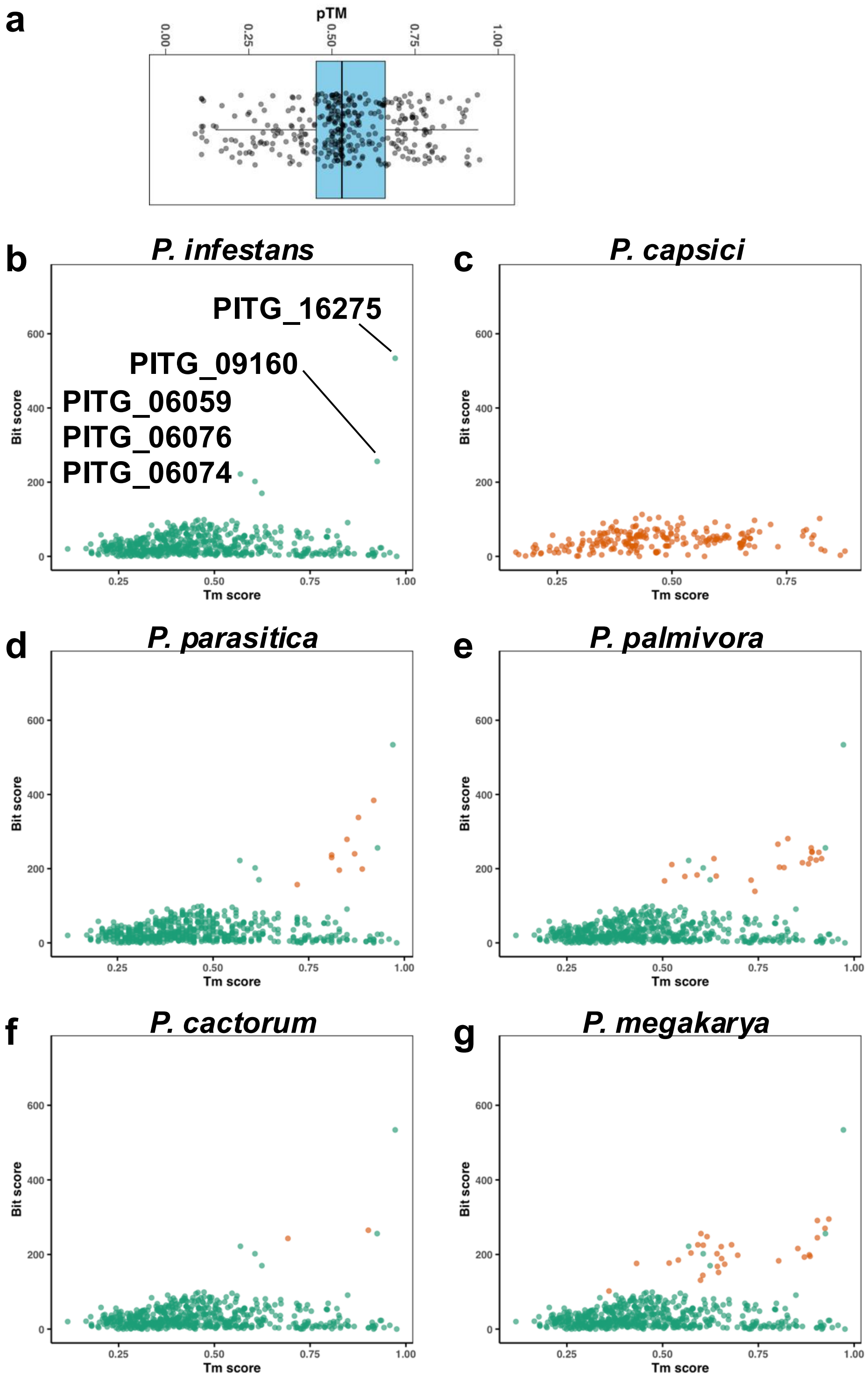


**Supplemental figure 17. Structural homologs of PITG_16275 are found in several *Phytophthora* species, but not *P. capsici*. (a)** Alphafold 3 was used to predict the structures of the RXLR effectors predicted from *P. capsici*. The highest-ranking model for each effector was chosen, a distribution of pTM scores for these effectors is shown. **(b)** *P. infestans* contains four additional effectors with predicted structural similarity. A Foldseek search was performed using a local database of *P. infestans* effector models (generated by Seong and Krasileva, 2023), a new model of PITG_16275 was used as an input. Each effector is represented by a data point in the plot. The hits with high Bit score and Tm score were inspected as possible structural homologs. **(c)** No *P. capsici* effector structures were identified with similarity to PITG_16275. A Folkseek search was performed using a structure of PITG_16275 predicted by Alphafold, against a library of *P. capsici* effectors whose structures were predicted using Alphafold 3. **(d)** Unlike *P. capsici*, structural homologs of PITG_16275 were identified in *P. parasitica*, as well as *P. palmivora* **(e)**, *P. cactorum* **(f)** and *P. megakarya* **(g)**. These hits are indicated in orange, their position within the plot from panel B is shown.

a

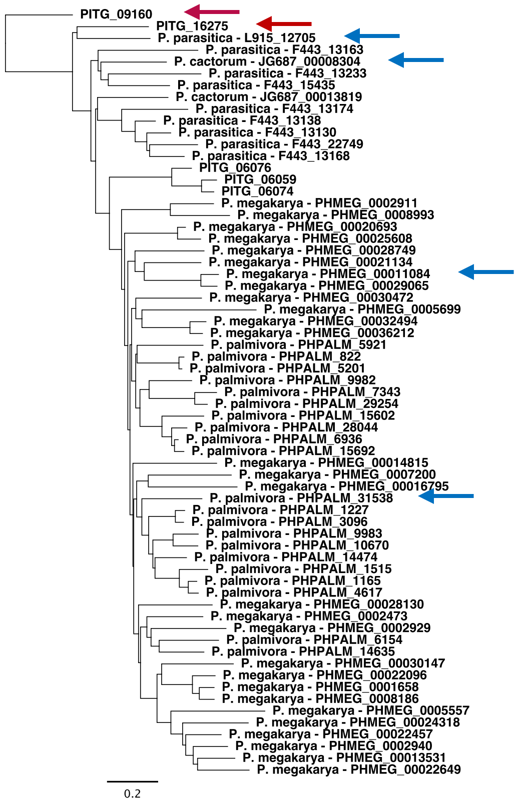


b

| | PITG_09160 | P. cactorum | P. megakar... | P. palmivora | PITG_16275 | P. parasitica |
|---|---|---|---|---|---|---|
| PITG_09160 | | 29.9% | 29.0% | 29.9% | 38.8% | 40.8% |
| P. cactorum | 29.9% | | 43.8% | 38.5% | 45.5% | 43.4% |
| P. megakarya | 29.0% | 43.8% | | 51.0% | 48.5% | 50.0% |
| P. palmivora | 29.9% | 38.5% | 51.0% | | 42.4% | 51.5% |
| PITG_16275 | 38.8% | 45.5% | 48.5% | 42.4% | | 60.0% |
| P. parasitica | 40.8% | 43.4% | 50.0% | 51.5% | 60.0% | |

c

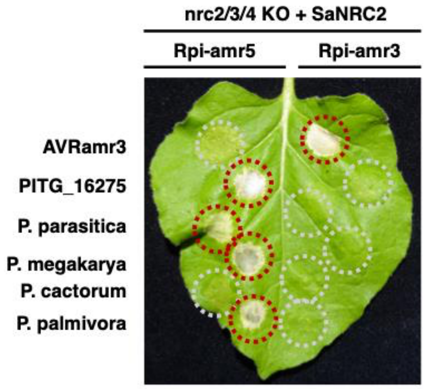


**Supplementary figure 18. Rpi-amr5 recognises PITG_16275 orthologs from *P. parasitica*, *P. megakarya* and *P. palmivora*. (a)** Orthologs of PITG_09160 and PITG_16275 are found in *P. megakarya* (28), *P. palmivora* (21), *P. parasitica* (8), and *P. cactorum* (2). Orthologs were identified using NCBI BLAST and local BLAST performed on the *Phytophthora* reference genomes. PITG_09160 and PITG_16275 are indicated with red arrows. Cloned orthologs from *P. megakarya*, *P. palmivora*, *P. parasitica,* and *P. cactorum* are indicated by blue arrows, these were selected as the highest scoring hits from the Foldseek search. **(b)** Heatmap of the recognised *P. infestans* effectors with the cloned orthologs from *P. megakarya*, *P. palmivora*, *P. parasitica* and *P. cactorum*. **(c)** Rpi-amr5 can recognise PITG_16275 orthologs from *P. megakarya*, *P. palmivora,* and *P. parasitica.* No HR was observed upon co-expression with the cloned *P. cactorum* ortholog. Effectors were co-expressed with sensor NLRs and SaNRC2 in the *N. benthamiana nrc2/3/4* KO line. *Agrobacterium* strains infiltrated at 0.3 $OD_{600}$, images were taken at 3 dpi.

**a**

| | P.cap_E144 | P.cap_E149 | P.cap_E150 | PITG_09160 | PITG_16275 |
|---|---|---|---|---|---|
| P.cap_E144 | | 65.4% | 65.6% | 10.4% | 15.2% |
| P.cap_E149 | 65.4% | | 92.5% | 11.8% | 15.9% |
| P.cap_E150 | 65.6% | 92.5% | | 11.8% | 15.2% |
| PITG_09160 | 10.4% | 11.8% | 11.8% | | 39.2% |
| PITG_16275 | 15.2% | 15.9% | 15.2% | 39.2% | |

**b**

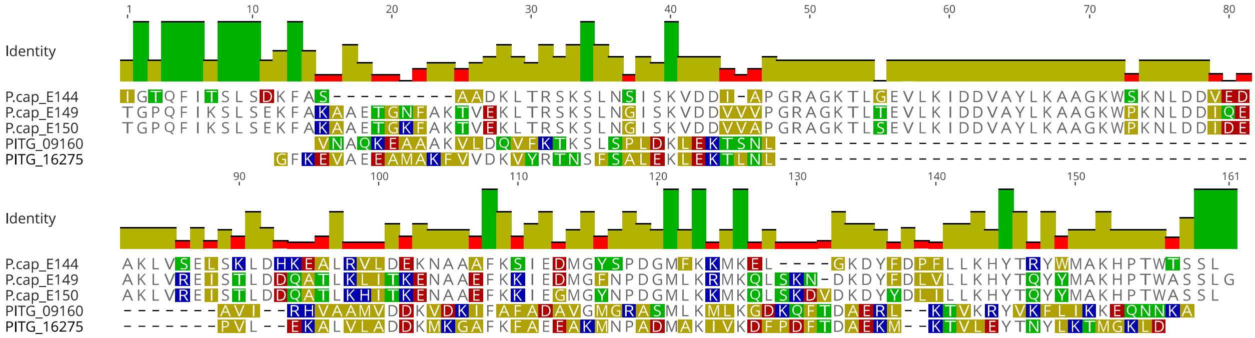


**Supplementary figure 19. Rpi-amr5 and Rpc2 recognise three *P. capsici* effectors with low sequence similarity to PITG_16275 or PITG_09160. (a)** Heatmap showing the percent identity of the post-EER amino acid sequences for the five effectors recognized by Rpi-amr5 and Rpc2. PITG_09160 and PITG_16275, from *P. infestans*, have low sequence similarity to PcE144, PcE149 and PcE150, from *P. capsici*. **(b)** Alignment of the five effectors (sequence after the EER motif) recognised by Rpi-amr5 and Rpc2.

**a**

| | PITG_16275 | PITG_09160 | |
|---|---|---|---|
| SP2275 | 2 | 2 | Complete Rpi-amr5 |
| SP2308 | 2 | 2 | |
| SP1102 | 2 | 2 | |
| SP2273 | 2 | 2 | |
| SP3400 | 2 | 2 | |
| SP2360 | 2 | 2 | |
| SP2297 | 2 | 2 | |
| SP1101 | 0 | 1.5 | Frameshift or premature stop in Rpi-amr5 |
| SP3409 | 0 | 1 | |
| SP2307 | 0 | 1.5 | |
| SP3406 | 0 | 0 | |
| SP1032 | 0 | 1 | |
| SP3399 | 0 | 2 | |
| SP2272 | 0 | 1.5 | |
| SP2298 | 0 | 1 | |
| SP2300 | 0 | 1 | |
| SP2271 | 0 | 2 | No Rpi-amr5 |

**b**

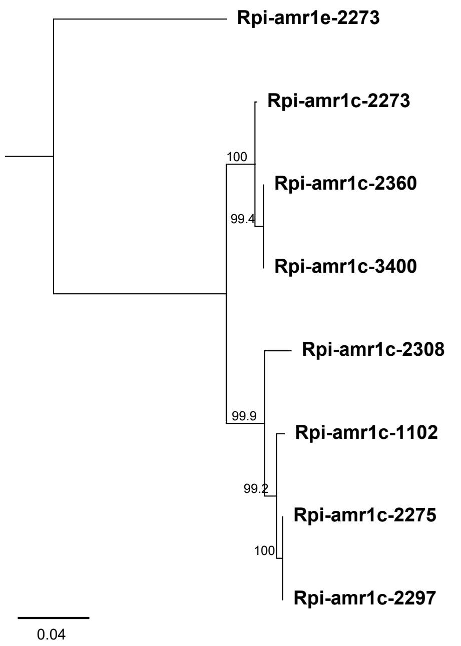


**c**

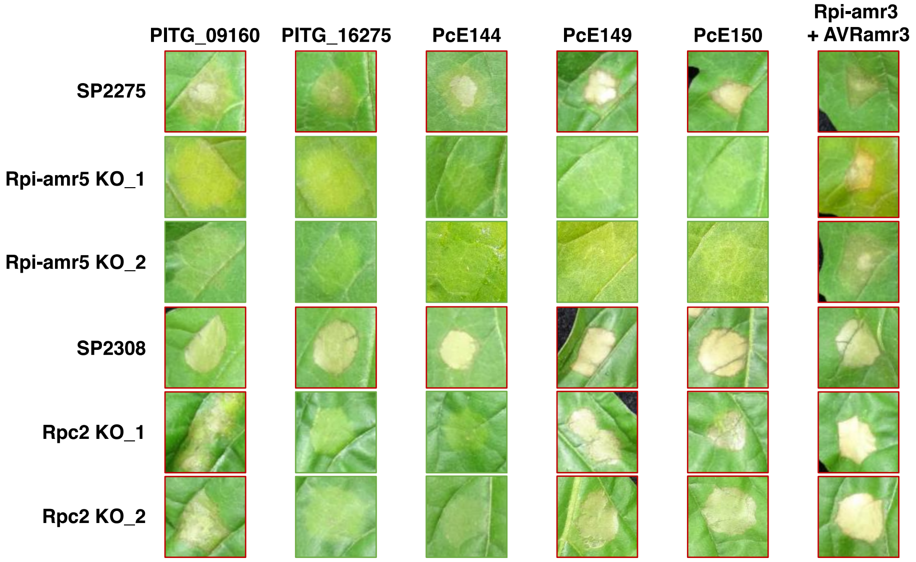


**Supplementary figure 20. PITG_09160 recognition is reported in several *S. americanum* accessions that lack complete *Rpi-amr5* alleles. (a)** Published PacBio RenSeq datasets were manually investigated for the presence of *Rpi-amr5* alleles, seven accessions were found to contain complete predicted alleles, 9 contained incomplete alleles with frameshift or premature stop mutations, for accession SP2271 no corresponding NLR was identified. PITG_16275 recognition occurs only when a complete *Rpi-amr5* allele is found, PITG_09160 recognition also occurs in accessions lacking a complete *Rpi-amr5* allele. HR scoring represented is taken from Lin et al 2023, 2 indicates strong cell death, 1 weak cell death, and 0 an absence of cell death. **(b)** Phylogenetic tree of Rpi-amr5 proteins predicted using *S. americanum* PacBio RenSeq datasets, constructed using complete protein sequences. the tree is rooted using Rpi-amr1-2273. **(c)** Agrobacterium infiltration of *Rpc2* knockout lines reveals that SP2308 contains an additional receptor capable of recognising PITG_09160, PcE149 and PcE150. Two independent biallelic mutants for both *Rpi-amr5* and *Rpc2* are shown.

**Supplementary table 1. Four NLR-encoding genes were identified as *Rpi-amr5* candidates.** *Rpi-amr5* maps to the *Rpi-amr1* cluster on Chromosome 11. Of the 16 paralogs identified in the SP2275 reference genome, four are candidates for *Rpi-amr5*. The ID, expression (determined using cDNA RenSeq of SP2275) and homology to the characterised *Rpi-amr1* cluster (from SP2273) is shown.

| **NLR_gene_ID** | **Expression in SP2275** | **Notes** | **Retained as a candidate?** |
|---|---|---|---|
| *NLR_1-2275* | Expressed | **Premature stop** | **-** |
| *NLR_2-2275* | **No expression** | - | **-** |
| *NLR_3-2275* | Expressed | - | **Yes** |
| *NLR_4-2275* | **No expression** | - | - |
| *NLR_5-2275* | **No expression** | - | - |
| *NLR_6-2275* | Expressed | - | **Yes** |
| *NLR_7-2275* | Expressed | **Frameshift mutation** | - |
| *NLR_8-2275* | Expressed | **Partial – No CC domain** | - |
| *NLR_9 (Rpi-amr1h-2275)* | Expressed | **Constitutive activity in *N. benthamiana*** | **Yes** |
| *NLR_10-2275* | Expressed | **Partial – No CC domain** | - |
| *NLR_11-2275* | Expressed | **Frameshift mutation** | - |
| *NLR_12-2275* | Expressed | **Identical to non-*Rpi-amr5* accession** | - |
| *NLR_13 (Rpi-amr1c-2275)* | Expressed | **Constitutive activity in *N. benthamiana*** | **Yes** |
| *NLR_14-2275* | Expressed | **Incomplete** | - |
| *NLR_15-2275* | Expressed | **Partial – incomplete NB-ARC, no LRRs** | - |
| *NLR_16-2275* | Expressed | **Identical to non-*Rpi-amr5* accession** | - |

**Supplementary table 2. Five NLR-encoding genes were identified as *Rpc2* candidates.** Using bulked segregant analysis, *Rpc2* was mapped to the *Rpi-amr1*/*Rpi-amr5* cluster on Chromosome 11. Of the 8 paralogs identified within the *Rpc2* mapping interval, five were retained as candidates. The ID and homology to the characterised *Rpi-amr1* cluster is shown.

| NLR_gene_ID | Candidate notes | Retained as a candidate? |
|---|---|---|
| *NLR_1-2308 (Rpi-amr1h-2308)* | - | **Yes** |
| *NLR_2-2308 (Rpi-amr1l-2308)* | - | **Yes** |
| *NLR_3-2308 (Rpi-amr1e-2308)* | - | **Yes** |
| *NLR_4-2308* | **Pseudogenised, 5 Kb insertion into NB-ARC** | - |
| *NLR_5-2308 (Rpi-amr1c-2308)* | **Constitutive activity in *N. benthamiana*** | **Yes** |
| *NLR_6-2308* | **Incomplete NB-ARC** | - |
| *NLR_7-2308 (Rpi-amr1a-2308)* | **Constitutive activity in *N. benthamiana*** | **Yes** |
| *NLR_8-2308* | **No clear ORF** | - |

**Supplementary table 3. Effectors recognised by Rpi-amr1e-2308, Rpi-amr5, or Rpc2.** *P. capsici* and *P. infestans* effectors were screened to identify elicitors of Rpi-amr1e-2308, Rpi-amr5 and Rpc2. Six effectors were identified in this study, five recognised by Rpi-amr5/Rpc2, and a *P. capsici* homolog of AVRamr1 (PcapAVRamr1) that is recognised by both Rpi-amr1e-2273 and Rpi-amr1e-2308. The ID (PITG for *P. infestans* effectors, PcE for *P. capsici* effectors), the gene which encodes the effector, and the amino acid sequence of each effector (after the EER motif) is shown below.

| Effector ID | Gene ID | Recognised by | Amino acid sequence post-EER |
|---|---|---|---|
| PITG_09160 | PITG_09160 | Rpi-amr5, Rpc2 | VNAQKEAAAKVLDQVFKTKSLSPLDK LEKTSNLAVIRHVAAMVDDKVDKIFAF ADAVGMGRASMLKMLKGDKQFTDAE RLKTVKRYVKFLIKKEQNNKA |
| PITG_16275 | PITG_16275 | Rpi-amr5, Rpc2 | GFKEVAEEAMAKFVVDKVYRTNSFSA LEKLEKTLNLPVLEKALVLADDKMKG AFKFAEEAKMNPADMAKIVKDFPDFT DAEKMKTVLEYTNYLKTMGKLD |
| PcE144 | DVH05_026132 | Rpi-amr5, Rpc2 | IGTQFITSLSDKFASAADKLTRSKSLNSI SKVDDIAPGRAGKTLGEVLKIDDVAYL KAAGKWSKNLDDVEDAKLVSELSKLD HKEALRVLDEKNAAAFKSIEDMGYSPD GMFKKMKELGKDYFDPFLLKHYTRYW MAKHPTWTSSL |
| PcE149 | DVH05_000363 | Rpi-amr5, Rpc2 | TGPQFIKSLSEKFAKAAETGNFAKTVEK LTRSKSLNGISKVDDVVVPGRAGKTLT EVLKIDDVAYLKAAGKWPKNLDDIQE AKLVREISTLDDQATLKLITKENAAEFK KIEDMGFNPDGMLKRMKQLSKNDKDY FDLVLLKHYTQYYMAKHPTWASSLG |
| PcE150 | DVH05_026158 | Rpi-amr5, Rpc2 | TGPQFIKSLSEKFAKAAETGKFAKTVEK LTRSKSLNGISKVDDVVAPGRAGKTLS EVLKIDDVAYLKAAGKWPKNLDDIDE AKLVREISTLDDQATLKHITKENAAEF KKIEGMGYNPDGMLKKMKQLSKDVD KDYYDLILLKHYTQYYMAKHPTWASS L |
| PcE173 (PcapAVRamr1) | DVH05_027339 | Rpi-amr1e-2308, Rpi-amr1e-2273 | LAGKNMFNAEKIEKALQDTSYAKTLFR RWKRYEVEHGAAFDKLIKFNIGKDDK VFGLYKSYVSWLEKHHPLGAETGGGP NLFSKAKLDKAMKDPKYENTMFGRW KRQGFESDAAYNKLLAFNLASDADVY KIYTKYVTWLNIHHPLAKTRKTTAKDF LFNVDRIARAKKDSEFAETLFKKWKTS GLDEKPVYLKLWDMGLKTDDELYKLY KNYVKWLDIHYPLPAKAT |

**Supplementary table 4. NLR-encoding genes characterized in this study.** Three novel NLR-encoding genes were characterized in this study - *Rpi-amr1e-2308*, *Rpi-amr5* and *Rpc2*. The gene names, reference sequence, and accession from which they were cloned is detailed below. Sequence references are as described by Lin et al 2023, and are available at https://figshare.com/projects/The_Solanum_americanum_pangenome_and_effectoromics_reveals_new_resistance_genes_against_potato_late_blight/145449

| ***R*-gene** | **Cloned from accession** | **Reference contig** | **Gene ID/position** |
|---|---|---|---|
| *Rpi-amr5* | SP2275 | utg376 | 754,620-759,628 |
| *Rpi-amr1e-2308* | SP2308 | Contig_24_SP2308_SMRT_RenSeq_RH_5_20__1 | nlr_3 |
| *Rpc2* | SP2308 | Contig_4_SP2308_SMRT_RenSeq_RH_5_20__1 | nlr_2 |
